# piTracer - Automatic reconstruction of molecular cascades for the identification of synergistic drug targets

**DOI:** 10.1101/2023.04.08.535933

**Authors:** Daniel Gomari, Iman W. Achkar, Elisa Benedetti, Jan Tabling, Anna Halama, Jan Krumsiek

**Author notes:** <u>Corresponding authors:</u> Anna Halama, Jan Krumsiek.

## Abstract

Cancer cells frequently undergo metabolic reprogramming as a mechanism of resistance against chemotherapeutic drugs. Metabolomic profiling provides a direct readout of metabolic changes and can thus be used to identify these tumor escape mechanisms. Here, we introduce piTracer, a computational tool that uses multi-scale molecular networks to identify potential combination therapies from pre- and post-treatment metabolomics data. We first demonstrate piTracer’s core ability to reconstruct cellular cascades by inspecting well-characterized molecular pathways and previously studied associations between genetic variants and metabolite levels. We then apply a new gene ranking algorithm on differential metabolomic profiles from human breast cancer cells after glutaminase inhibition. Four of the automatically identified gene targets were experimentally tested by simultaneous inhibition of the respective targets and glutaminase. Of these combination treatments, two were be confirmed to induce synthetic lethality in the cell line. In summary, piTracer integrates the molecular monitoring of escape mechanisms into comprehensive pathway networks to accelerate drug target identification. The tool is open source and can be accessed at https://github.com/krumsieklab/pitracer.

## 1 Introduction

Drug-resistance is one of the major challenges of cancer treatment and is estimated to cause up to 90% of cancer-related deaths^1,2^. Genetic, epigenetic and microenvironmental modifications promote the tumor’s intrinsic functional plasticity, which enables the development of cellular *escape mechanisms* that allow the cells to overcome treatment effects. A wide variety of such escape mechanisms has been described in the literature^2^, among which the reprogramming of cellular metabolic processes has recently emerged as a fundamental element in the development of resistance^3–6^. There is increasing evidence that cancer cells, under drug-imposed selective pressure, can rewire their metabolic processes towards alternative pathways^7–10^. To identify effective therapy options for tumors that have undergone metabolic reprogramming, it is essential to characterize the specific metabolic escape mechanisms that cancer cells are exploiting.

Metabolomics measurements provide an ideal readout for capturing metabolic reprogramming mechanisms in a tumor, since metabolite levels represent the current state of cellular regulatory processes and energy dynamics^11,12^. Several previous studies have already showcased how metabolomic profiling can identify targetable escape mechanisms in cancer. For instance, Cardaci *et al*. utilized metabolomics to identify pyruvate carboxylase as an essential enzyme in succinate dehydrogenase (SDH)-deficient cancer cells, outlining a treatment strategy for SDH-associated malignancies^13^. Joly *et al*. demonstrated a synthetic lethal relationship between glucose transport and glutathione synthesis using metabolomics in cancer cells^14^. Moreover, we have previously used metabolic profiling to rationally identify molecular targets for combined treatment in breast cancer and lymphoma cell lines, respectively^15,16^. However, to identify the proposed therapeutic targets, all above-mentioned studies had to rely on a laborious and meticulous manual process of contextualizing statistical findings in terms of regulatory cascades and biochemical pathways.

To facilitate this process, we developed *piTracer*, a computational framework to automatically identify targetable escape mechanisms based on measured metabolomic changes. The method builds on an extensive, integrated and manually refined gene-regulatory and metabolic network, which is combined with a novel path-finding and gene scoring algorithm for target prioritization. Its core feature is the ability to traverse the integrated network and automatically identify biologically valid cascades between two molecules, genes or metabolites, highlighting various alternative paths through the network. Based on these cascades, piTracer then identifies potential metabolic escape mechanisms and ranks genes by their potential as therapeutic targets to block those escape mechanisms. We refer to these combinations as *synthetic lethal* pairs; notably this terminology is commonly used in literature for combinations of gene mutations^17^, but also for combinations of gene knockout interventions that lead to cell death^18^.

A variety of other methods leveraging omics profiles to computationally identify drug targets and synthetic lethal combinations for cancer therapy have previously been published. For example, several studies screened large sets of target combinations using genome-scale metabolic models followed by experimental validation^19,20^. Lu *et al*. used metabolomics profiles to gain insights into the synergistic action of two drugs after a high-throughput combination screening^21^. Ruppin and colleagues leveraged large-scale genomics and RNA sequencing data to generate statistical evidence for synthetic lethality, leading to the prediction of novel combination therapy that were retrospectively tested in clinical trials^22–24^. The MAPPS framework from Riaz et al. provides a drug target identification functionality based on KEGG metabolic networks based on a predefined lists of source and target metabolite lists^25^.

The unique feature of piTracer lies in its core prediction paradigm: It leverages the metabolic alterations resulting from single-drug pre- vs. post-treatment experiments to computationally identify the underlying escape mechanism of the tumor cell. This allows to automatically identify genes with a high likelihood of inducing synthetic lethality when targeted with a second drug, thereby gaining insights into tumor resistance mechanisms and drastically reducing the number of validation experiments to be performed.

In this paper, we first introduce the piTracer framework and validate its ability to retrieve molecular cascades by reconstructing the enzymatic steps involved in well-known metabolic pathways using only the substrate and the final product of the pathways. Moreover, we reconstruct gene/metabolite cascades reported in a metabolomics genome-wide association study (mGWAS), which study the effects of naturally occurring genetic variants on the metabolic network. Finally, we apply the drug target prediction algorithm on metabolomics data from a breast cancer cell line treated with a glutaminolysis inhibitor and then experimentally validate selected targets.

## 2 Results

### 2.1 The piTracer framework

The piTracer framework enables the accurate and automatic reconstruction of biological cascades. Its backend consists of a combined gene regulatory and metabolic network, constructed from Recon 3D^26^, STRING v11^27^, the OmniPath database^28^, and a gene regulatory network created by Sonawane *et al*.^29^ based on data from STRING and the Genotype-Tissue Expression (GTEx) project^30^. Each metabolite can appear multiple times in the network depending on its cellular localization (referred to as “compartment”), e.g., extracellular, cytosol, mitochondrion, etc. The network integration process included extensive curation of the primary data sources and the application of a new atom tracing-based processing method that removes shortcut nodes such as metabolic cofactors (see Methods). The final network consisted of 7,412 unique genes and 3,143 unique metabolites, with 4,631,052 directed interactions between them (**Figure 1a**). Based on this network, we developed a shortest path-based graph algorithm that enumerates biological cascades between any two entities in our multi-layered network, i.e., two metabolites, two genes, or a gene and a metabolite (example shown in **Figure 1b**). We then developed an approach to predict potential therapeutic targets based on the proximity of genes to the differential metabolites from a drug treatment experiment (**Figure 1c**).

**Figure 1.**
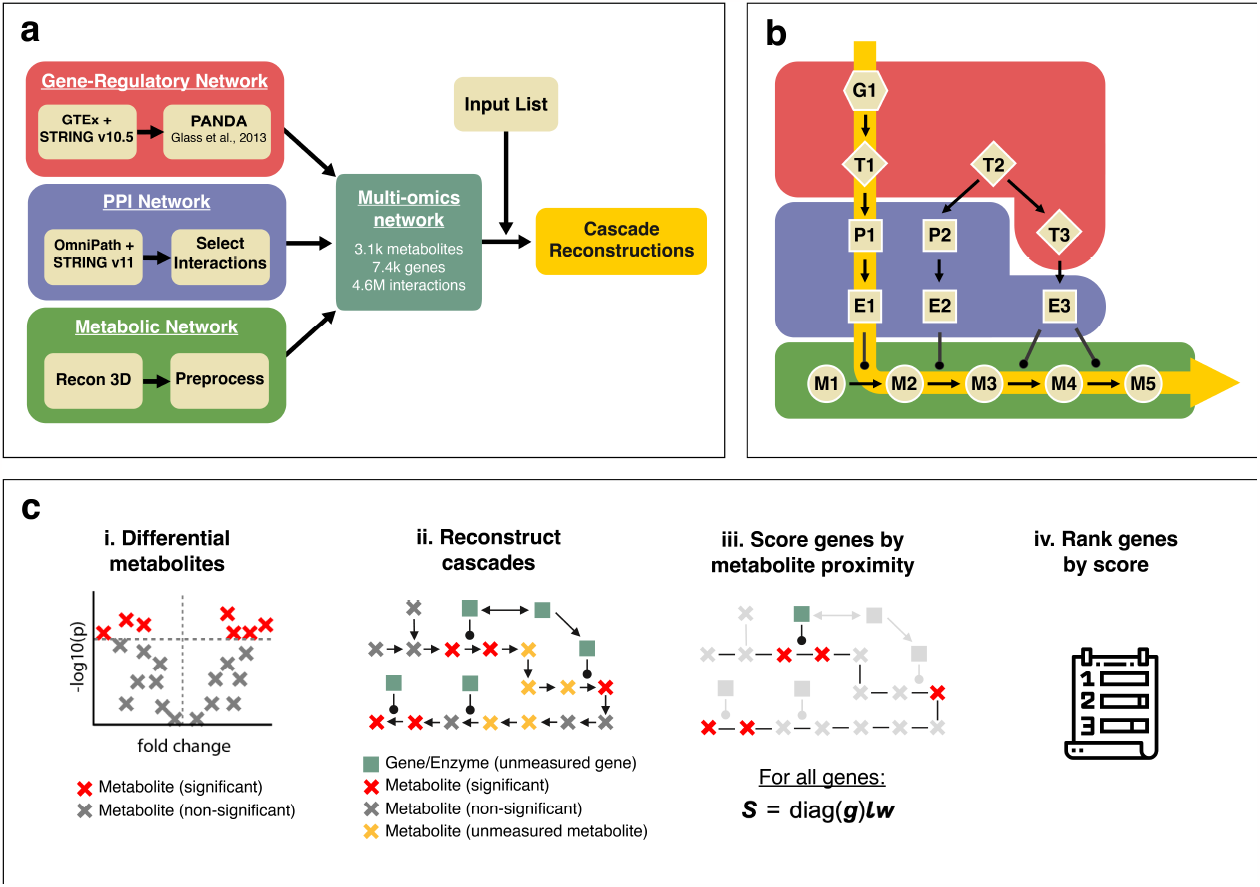
piTracer framework. **(a)** The piTracer backend. The multi-omics network is constructed from gene-regulatory edges, a protein-protein interaction (PPI) network, and a curated version of the Recon3D genome-wide human metabolic reconstruction. The input to the reconstruction method is a list of genes or metabolites, e.g., derived from the statistical analysis of an experiment, and the output are reconstructed biological cascades between these molecules. **(b)** Example of a reconstructed, shortest path-based cascade between a transcription factor gene G1 and its downstream effect on metabolite M5 in a metabolic pathway. G: gene, T: transcription factor, P: protein, E: enzyme, M: metabolite. **(c)** Overview of the gene target scoring algorithm. (i) An input list of differential metabolites is supplied to the algorithm. (ii) All gene regulatory and metabolic steps connecting the metabolites, together with the enzymes involved in each reaction, are reconstructed. (iii) and (iv) the genes involved in the reconstructions are then scored and ranked based on their proximity to significantly affected metabolites.

All piTracer functionalities are available as R scripts at https://github.com/krumsieklab/pitracer, which also links to a hosted, interactive Shin application. As input, the app accepts HGNC gene symbol^31^ identifiers for genes, and HMDB^32^, KEGG^33^, PubChem^34^, or Recon 3D^26^ identifiers for metabolites. The online application provides interactive visualizations of reconstructed molecular cascades and a downloadable list of predicted gene targets.

### 2.2 Validation of piTracer core functionality by reconstructing known biological pathways

To validate piTracer’s ability to find correct paths between molecule pairs, we first reconstructed the steps from a series of known biochemical cascades. This included several manually selected, well-established metabolic synthesis reactions involved in central carbon metabolism, gene-to-metabolite associations with known reaction cascades based on a metabolomics genome-wide association studies (GWAS) study^35^, and gene-to-gene immune response and developmental signaling cascades. A schematic of the type of reconstruction results produced by piTracer is shown in **Figure 2a**. For comparison, all reconstructions were also performed using two well-known computational tools that allow for metabolite-metabolite, gene-metabolite, and gene-gene tracing: Ingenuity Pathyway analysis (IPA)^36^ and ConsensusPathDB (CPDB)^37^.

**Figure 2.**
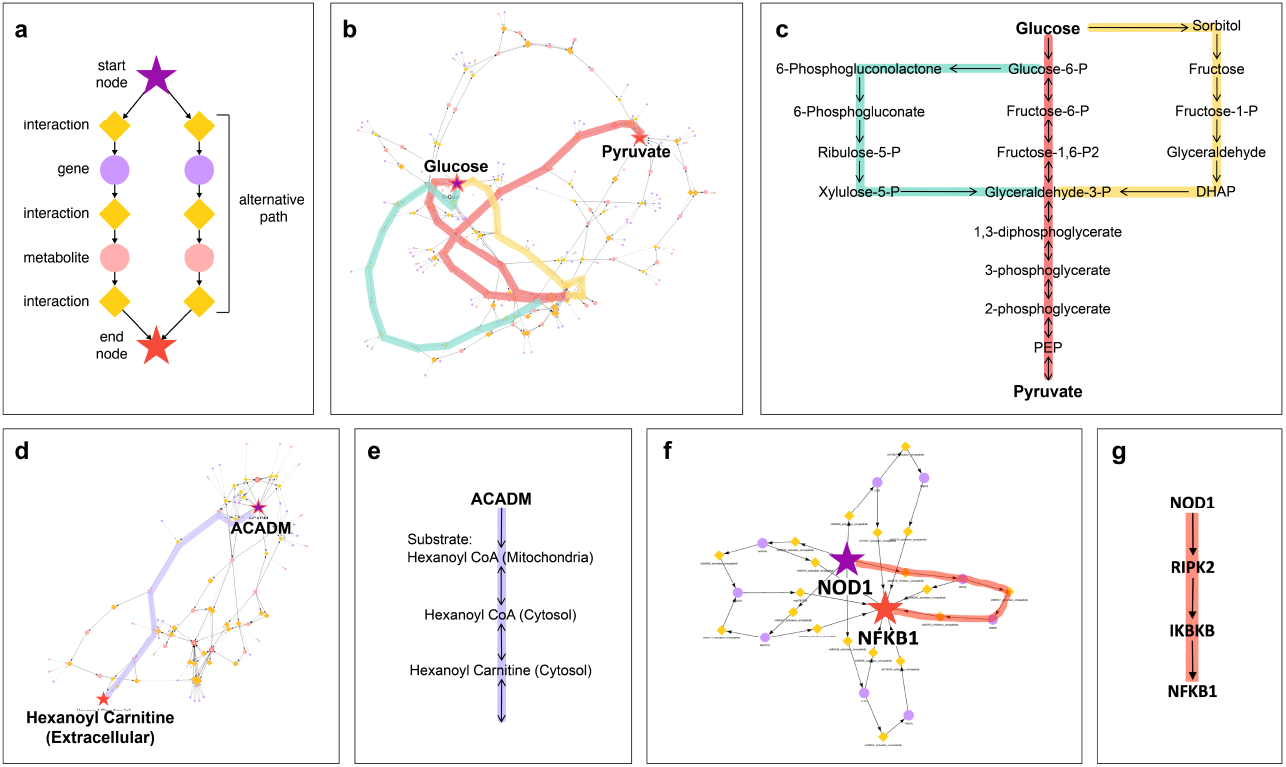
piTracer validation on known metabolic pathways. **(a)** Schematic of reconstruction results produced by piTracer. For each start node to end node pair, multiple possible paths are illustrated. Connections between two metabolites/genes are encoded in “interaction” nodes, which contain biological function such as “activates” or “catalyzes”. **(b)** Screenshot of the glycolysis, pentose phosphate, and sorbitol pathways as generated by the piTracer app. Cascades were reconstructed using glucose as the starting node and pyruvate as the end node. Three well-known carbohydrate metabolic pathways were manually highlighted. Note that for the pentose phosphate pathway, not all branched routes were highlighted for the visualization, but are present in the reconstruction. **(c)** Textbook versions of the glycolysis, pentose phosphate, and sorbitol pathways, which are identical to the piTracer reconstructions in (a). P: phosphate, DHAP: dihydroxyacetone phosphate, PEP: phosphoenol pyruvate, CoA: coenzyme A. **(d)** piTracer reconstruction between ACADM and hexanoyl carnitine. **(e)** The carnitine shuttle pathway and extracellular transport of acylcarnitines, which corresponds to the piTracer path in (d). **(f)** piTracer reconstruction of the pathway between NOD1 and NFKB1, and **(g**) the corresponding literature-based version of the pathway.

We started by reconstructing the enzymatic steps of glucose catabolism by connecting glucose and pyruvate. The algorithm produced three well-known pathways: The glycolysis pathway, the pentose phosphate pathway, and the sorbitol pathway (**Figure 2b+c**). Other paths in the reconstructed cascade also represent valid molecular steps, such as inositol metabolism, phospholipid biosynthesis, and the conversion of serine to pyruvate through serine dehydratase (SDS)^38^. Comparing our results to the reconstructions from IPA and CPDB, neither method was able to reconstruct these central carbon metabolic pathways at the same resolution as piTracer (**Supplementary Figure 1**). We furthermore reconstructed the steps of the citric acid cycle using citrate and oxaloacetate as inputs (**Supplementary Figure 2**), where piTracer correctly recreated all individual reaction steps, while neither IPA nor CPDB properly reconstructed the pathway. Notably, IPA did additionally show a correct version of the TCA cycle. However, it was a graphical representation of literature-curated version of the pathway, while we here focus on the unbiased reconstruction of pathways directly from the underlying networks.

The second set of validations was performed by reconstructing molecular cascades reported in a metabolomics GWAS by Shin et al^35^. Genetic variation has been shown to affect blood metabolite levels, and these associations are often reflective of known biochemical reactions^35,39^. We first investigated the association between a SNP in the ACADM gene and blood hexanoylcarnitine. Briefly, ACADM is an enzyme involved in mitochondrial beta-oxidation and hexanoylcarnitine is an acylcarnitine, which is a transport form of one of the substrates of this enzyme^40^. The genetic link between ACADM SNPs and hexanoylcarnitine has been described and replicated in multiple studies^35,41,42^. piTracer was able to correctly reconstruct the steps in this molecular cascade, including the ACADM reaction^43^, the mitochondrial transport of CoA-carnitine to the cytosol, as well as the transport from the cytosol to the extracellular space (**Figure 2d+e**). Arguably, the extracellular space is the best representative compartment for circulating blood metabolites in our context. IPA and CPDB again produced misleading or incomplete reconstructions (**Supplementary Figure 1**).

As an interesting example of complex biochemical reactions that are still under active investigation, piTracer correctly identified that the association between SNP rs2403254 and 2-hydroxyisovalerate was driven by the LDHA gene, which is located at the genetic locus of that SNP (**Supplementary Figure 5**). The original GWAS publication had incorrectly annotated this association with HPS5^35^, the gene that contains the SNP in an intron. This finding was later refuted, and the SNP-metabolite association was experimentally shown to be caused by LDHA^44^. As further examples, piTracer correctly reconstructed pathway cascades between ALDH18A1 and citrulline as well as MCCC1 and 3-hydroxyisovalerylcarnitine (**Supplementary Figures 3&4**, respectively) from the Shin *et al*. GWAS^35^.

The reconstruction capabilities of gene-to-gene signaling cascades were validated on steps of the NOD-like receptor signaling pathway using NOD1 and NFKB1 (**Figure 2f+g** and **Supplementary Figure 6**), the NODAL signaling pathway utilizing NODAL and its inhibitor LEFTY1, and the Hippo signaling pathway with STK3 and TEAD1 (**Supplementary Figure 6**). In all cases, piTracer reconstructed the correct molecular paths, while IPA and CPDB either generated overly dense networks, or traces that did not include the complete set of molecular steps in the cascade.

### 2.3 Identification of synthetic lethal candidates for glutaminase inhibition in triple-negative breast cancer cells

In the next step, we evaluated the drug ranking algorithm of the piTracer framework. The method is built on the same background network and tracing techniques described in the previous sections. The case study is based on data from a previously published analysis, where we showed that suppressing glutaminolysis with the glutaminase (GLS) inhibitor C968 is insufficient to induce cancer cell death in the glutamine-dependent triple negative breast cancer cell line MDA-MB-231^16^. Using metabolic profiling, the study demonstrated that this survival is due to the activation of lipid catabolism and autophagy, which arise as escape mechanisms from glutaminolysis inhibition.

We used our computational framework to rank genes by their potential for synthetic lethality based on the list of significantly up- and down-regulated metabolites from that experiment (**Supplementary Table 1)**. Briefly, a target score was calculated for each gene based on their proximity to significant metabolites, penalized by the number of enzymes the gene can reach overall, which reduces false positives that might arise due to hub transcription factors affecting the majority of the network (see detailed description in the **Methods**). Using this approach, we ranked a total of 7,253 genes by their potential to cause synthetic lethality in MDA-MB-231 cells treated with glutaminase inhibitor (C968). The full list of gene ranks can be found in **Supplementary Table 2**.

The highest-scoring gene was CPT1 (isoforms CPT1A, CPT1B, and CPT1C), which has been demonstrated to induce synthetic lethality with glutaminase inhibitors (C968 and CB-839) in our previous work^16^ and an independent study by other researchers^45^. This finding can be considered as an initial validation of the piTracer drug target prediction framework. We then selected 4 additional gene targets among the top 30 candidates from the gene list, based on their involvement in distinct metabolic pathways, druggability, and drug availability (Table 1). The selected targets for validation were: (1) PLD2, a phospholipase involved in lipid metabolism, which catalyzes the hydrolysis of phosphatidylcholine to produce phosphatidic acid and choline^46^. PLD2 was targeted by CAY01594. (2) SLC25A20, a carnitine-acylcarnitine translocase (CACT) essential for fatty acid oxidation^47^, which was targeted by Ingenol Mebutate. (3) AKR1A1, an aldo-keto reductase, which are detoxifying enzymes involved in the reduction of biogenic and xenobiotic aldehydes^48^. AKR1A1 was targeted by Imirestat. (4) AKR1B1, an aldo-keto reductase involved in the glucose-transforming polyol pathway (PP), which has recently been linked with the epithelial-to-mesenchymal transition^49^. AKR1B1 was targeted by Ranirestat.

**Table 1.**
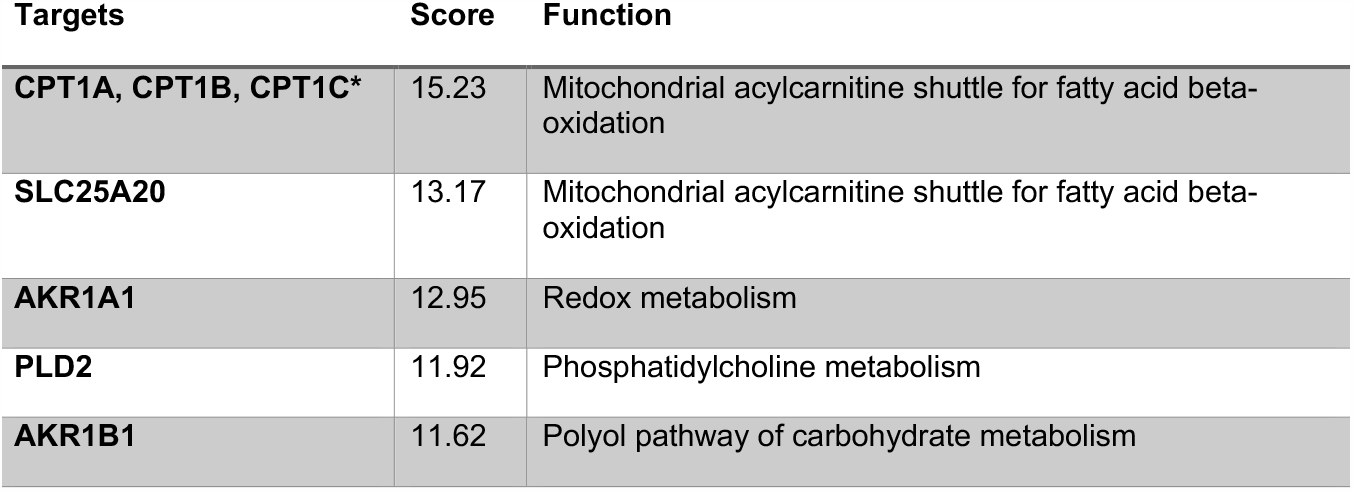
Targets for combination treatment with glutaminase inhibition. The genes were chosen from the top 30 list based on their involvement in different metabolic pathways and their potential to be targeted by drugs. *CPT1 inhibition has already been validated and showed to induce synthetic lethality with glutaminase inhibition in independent previous studies^16,45^.

### 2.4 Validation of synthetic lethal candidates under glutaminase inhibition

We next evaluated whether synthetic lethality can be achieved by simultaneously targeting GLS and the protein products of the four selected genes (AKR1A1, AKR1B1, PLD2, SLC25A20) in the MDA-MB-231 cell line. Before performing the cell viability experiments, we first confirmed that the cell line expressed the four proteins and that GLS inhibition with 10 μM of C968 did not affect the respective protein levels (**Supplementary Figure 7)**.

We then tested whether single-agent inhibition of any of the four proteins individually affected the viability of MDA-MB-231 cells. The cells were treated with the four selected drugs at five different concentrations and three time points (24h, 48h and 72h, **Supplementary Figure 8**). Overall, cell viability was not significantly affected by any of the tested treatment strategies, except for the inhibition of SLC25A20 at the highest concentration (100 μM) of Ingenol Mebutate, where we observed a 40% decrease in cell viability (**Supplementary Figure 8**). This indicates that AKR1A1, AKR1B1, and PLD2 are not regulating pathways that are crucial for the viability of MDA-MB-231 cells under standard conditions. However, the decrease in viability after inhibition of SLC25A20, a transporter involved in fatty acid oxidation, suggests that this process is of high metabolic relevance for these cells.

Next, we investigated whether simultaneous inhibition of glutaminase and each of the four piTracer targets led to significant decrease in viability of MDA-MB-231 cells beyond GLS inhibition. Cells were treated with two selected concentrations of the four drugs in combination with the GLS inhibitor C968 (10 uM) at different time points. We observed a time- and dose-dependent decrease in cell viability across all examined drugs (**Figure 3A**). In particular, the simultaneous inhibition of GLS and PLD2 (involved in glycerophospholipid metabolism) as well as GLS and SLC25A20 (involved in fatty acid metabolism) resulted in significant decreases in cell viability. The strongest effects were observed at 72h for GLS/SLC25A20 inhibition at the highest concentration (100 μM), where less than 20% of cells remained viable, and for GLS/PLD2 inhibition at the highest concentration (10 μM), where around 25% of cells remained viable. Notably, the inhibition of GLS and either AKR1A1 (aliphatic aldehyde metabolism) or AKR1B1 (polyol pathway of carbohydrate metabolism) did not lead to strong synergistic effects. The viability findings were further supported by microscopy images of the MDA-MB-231 obtained under the same conditions (**Supplementary Figure 9**).

**Figure 3.**
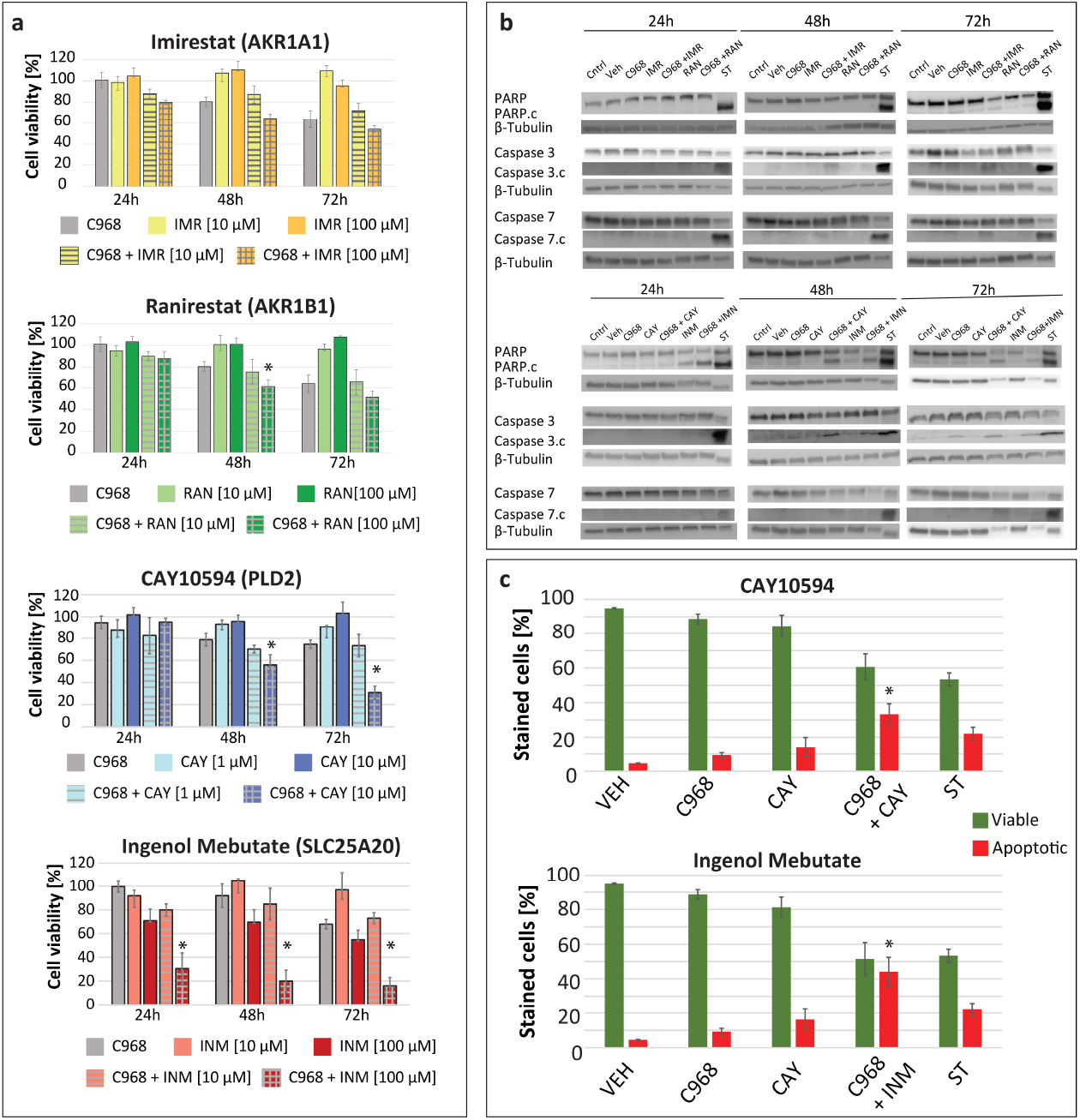
Combined treatment strategies determined by piTracer accelerate MDA-MB-231 cancer cell death. **(a)** Cell viability after treatment determined by MTT assays. A significant decrease in viability of the cells was observed for the combined inhibition of glutaminase and either PLD2 or SLC25A20 (**b)** Western blot analysis for apoptotic markers PARP & cleaved PARP, Caspase-3 & cleaved Caspase-3, and Caspase-7 & cleaved Caspase-7 demonstrated apoptosis in cells under combined inhibition of glutaminase and either PLD2 or SLC25A20. **(c)** Apoptosis was confirmed after cell staining with fluorescein-conjugated annexin-V and propidium iodide (PI) and analysis with flow cytometry. The percent distribution of live (green) and apoptotic cells (red) is presented. The cells assayed with Western blots and FACS were conditioned as follows: Cntrl - untreated; Veh - treated with DMSO; C968 - treated with 10 μM glutaminase inhibitor; CAY - treated with 10 μM PLD2 inhibitor (CAY10594); INM - treated with 100 μM SLC25A20 inhibitor (Ingenol Mebutate). IMR - Imirestat; RAN - Ranirestat; ST - staurosporine used as positive control (apoptosis inducer). *Indicates combined treatment with statistically significantly stronger effects compared to vehicle as well as to the two single treatments (C968 and selected agent).

The decrease in cell viability observed after combined targeting of glutaminolysis and PLD2 as well as SLC25A20 suggests the activation of apoptosis. Confirming this hypothesis, we observed an increased expression of the apoptotic markers PARP, Caspase-3, and Caspase-7 under simultaneous inhibition of GLS and either SLC25A20 or PLD2 (**Figure 3B**). Consistent with the results of the viability experiments, the apoptotic response was not observed in cells in which AKR1A1 or AKR1B1 were inhibited with GLS. We further validated the activation of apoptosis for the two effective drug combinations using fluorescence-activated cell sorting (FACS), where we observed a significantly higher number of apoptotic cells in those combined treatment groups compared to the single agent conditions (**Figure 3C**).

Taken together, these results show that piTracer correctly identified two MDA-MB-231 escape mechanisms under glutaminolysis inhibition and predicted key molecules in different metabolic pathways contributing to cancer cell survival.

## 3 Discussion

The piTracer framework provides a novel computational approach for the reconstruction of molecular pathways with a wide area of applications. The method can be used to rank of combination treatment candidates, based on the mapping of differential metabolomic profiles after drug treatment onto a highly curated molecular network. The central idea of piTracer is to identify the metabolic component of escape mechanisms in response to a drug, so that a second drug can be rationally chosen to induce a synergistic effect leading to cell death. This process significantly streamlines the testing process, allowing for the selection of only a limited, highly promising set of drug combinations. This carries significant potential for the drug development field, by reducing the time and resources required to identify lead combination candidates for further testing.

In this paper, we first performed a series of reconstruction steps on known pathways as a positive control step, and then tested the algorithm on a triple negative breast cancer cell line treated with the glutaminase inhibitor C968. Out of the five top-ranking combination targets that we considered, three were confirmed to have a synergistic effect with C968 (one from literature, two from our own validation experiments).

In addition to the drug target identification functionality, piTracer’s backend software provides a convenient tool for scientists to explore the relationships between molecules in a multi-omics fashion. To facilitate this exploratory process, we published piTracer’s underlying R code and made its functionalities available through a web portal, where users can interactively investigate the molecular paths connecting any two molecular entities (e.g., a metabolite and a gene transcript). This framework can be regarded as a “Google Maps” for cellular processes, highlighting alternative routes, path lengths, dead ends, and roadblocks on the way from point A to point B. This toolbox can prove valuable in many concrete applications, for example in gene-knockout experiments with differential metabolomics profiles, where piTracer can provide plausible biological mechanisms to explain how a gene of interest connects to the observed changes in metabolite levels. Future versions of the piTracer framework will integrate additional omics layers, such as transcriptomic and mutational profiling, which will add further levels of information for better path prioritization.

The ultimate goal of piTracer is the integration into real-world drug development pipelines, including classical discovery approaches for the identification of new pharmaceutical interventions. To this end, novel combination treatment strategies originating from the piTracer approach, such as the combined inhibition of glutaminase and PLD2 or SLC25A20 for triple-negative breast cancer discovered in this paper, should be further validated with established approaches and clinical trials to evaluate real clinical benefits. Moreover, our approach could be integrated into individualized precision medicine applications in the future. For example, patient-derived tumor organoids have recently gained attention as personalized models of a patient’s cancer^50^. Performing *in vitro* treatment experiments on organoids resistant to primary chemotherapy, fueled by metabolomics measurements and piTracer, has the potential to generate novel, individualized secondary treatment options for patients with refractory disease. A first successful study on treatment decisions informed by organoid-based experiments has recently been published in the context of breast cancer^51^. In our view, such individualized treatment approaches combined with the power of omics profiling and computational drug prioritization carries huge potential for a new era of precision oncology.

## 4 Online Methods

### 4.1 Metabolic network construction

Recon 3D^26^ constitutes the basis for our metabolic network. We performed the following preprocessing steps to ensure that metabolites were correctly connected and that shortcut nodes, such as cofactors, were pruned from the network.

We downloaded “RXN” reaction files from the Virtual Metabolic Human database (VMH)^52^. These files contain structural data for the reactants and products of each reaction, including information on atoms that are transferred from reactants to products. For metabolic reactions that did not have an RXN file in VMH, we generated them from MOL files downloaded from VMH (which define the molecular structure of each molecule) using the Reaction Decoder Tool v1.5.1^53^. To connect metabolites in the metabolic network, we created a special scoring system to ensure that for each step in the network there was a certain amount of atomic overlap from one molecule to the next. Briefly, we connected a reactant and a product of a reaction in our network if both shared a certain percentage of their atoms, while avoiding the creation of “biologically unreasonable” shortcuts between molecules that do not share a sufficient number of atoms. The detailed procedure is described in Supplementary Note 1. Despite careful design of this approach, manual inspection showed that reactions involving the binding of coenzyme A (CoA) to a metabolite were filtered out using our scoring method. For instance, the succinyl-CoA to succinate reaction, an important step in the tricarboxylic acid (TCA) cycle, was omitted from our metabolic network. Thus, we manually curated a list of these reactions and added them back into our metabolic network.

To further exclude cofactors from our metabolic network, we used a curated “blacklist” that includes nicotinamide adenine dinucleotide (NAD) and derivatives, flavin adenine dinucleotide (FAD) and derivatives, nucleotides, ions, CoA, acetyl CoA, and others. The blacklist can be found in **Supplementary Table 3**.

### 4.2 Gene regulatory network construction

We constructed a gene interaction network using the following databases: (1) STRING v11^27^ was used for the protein-protein interaction (PPI) network. Only PPIs containing directional information, i.e., activation, inhibition, and catalysis and with scores above 400 for the STRING “experiments” and “experiments transferred” scores were selected. (2) Signaling cascades were imported from the OmniPath database^28^ and interactions with OmniPath “consensus directionality” criteria were chosen. (3) Further gene regulatory information was imported into the multi-omics network by including a network created by Sonawane *et al*.^29^ using PANDA^54^ on data from the Genotype-Tissue Expression (GTEx) project^30^ and STRING without further preprocessing.

### 4.3 Cascade reconstructions

In the following, we describe the different algorithmic steps to extract paths and traces between a start and an end node from the multi-omics network.

#### k-shortest paths

Yen’s k-shortest path algorithm calculates the *k*-shortest paths between a pair of nodes in a network^55^. We used the implementation from the yenpathy R package (https://github.com/ecohealthalliance/yenpathy).

#### Path clustering

To improve the visualization of the *k*-shortest paths in a trace, especially for large values of *k*, we developed a clustering algorithm to group similar paths between the two molecules. We first compute the distances between the *k*-shortest paths between two nodes using the CoMapPa2 algorithm^56^. Briefly, CoMapPa2 calculates the similarity between two paths based on the number of steps required in the background network for each node in one path to reach all other nodes in the other path. Given these distances, a path dendrogram is then created using the hclust function in R. Path clusters are obtained through dynamic tree cutting using the cutreeHybrid function from the Dynamic Tree Cut R package^57^.

#### Heuristics for speed-up

To speed up the trace calculations, piTracer dynamically reduces the search space of the *k*-shortest paths algorithm per trace. This is achieved by confining the search to a subnetwork in the dense multi-omics network for each pair of start and end nodes. This subnetwork will contain a subset of all the shortest paths that exist between a pair of nodes. That is, the subnetwork might not contain all possible *k*-shortest paths for a trace and will cap the value of *k*, but this will significantly accelerate the calculation of traces. We implemented the heuristic as follows: (1) Initially, we extract the set *N*_*start*_ of all reachable nodes from the start node and another set *N*_*end*_ of all nodes that can reach the end node in a directed manner. (2) We define a hyperparameter *v*, such that we can calculate an overlap set *O* ⊆ (*N*_*start*_ ∩ *N*_*end*_) for which |*O*| ≤ *v*. For instance, *v* = 5 means that there are at most 5 molecules in *O*. Given that |*O*| ≤ *v*, the hyperparameter *v* determines the number of maximum shortest path *k* that can be calculated for a trace. We currently set *v* = 5, which on average allows 300 shortest paths to be found on our network, depending on the start and end nodes chosen. Increasing *v* increases the number *k*-shortest paths that can be calculated per trace in exchange for an increase in computational time. (3) We calculate a set *D*_*s*_ of the shortest distances between all pairs of the start node and nodes in *O*. We then extract the *v* shortest distances from *D*_*s*_ and select the maximum value *d*_*s*_. The same procedure is performed using the end node to calculate *d*_*e*_. (4) Afterwards, a subnetwork of the multi-omics network is created by including all nodes and edges within distance d_s_ towards the *O* nodes and distance d_e_ from the *O* nodes. (5) Tracing is then performed on the final, restricted multi-omics subnetwork.

### 4.4 Gene target prioritization

We prioritize gene targets based on metabolic changes after treatment with a primary drug using the following steps: (1) Perform standard differential abundance analysis on the metabolomic dataset before and after treatment with a single drug. (2) Use piTracer to calculate the 10-shortest paths between all metabolite pairs in the dataset to create a “context network” called ***C***. (3) Assign weights ***w*** to metabolites in ***C***, with ***w***=1 for significantly altered metabolites and ***w***=0 for the rest, which includes both non-significant metabolites and additional metabolites included in ***C*** that are not present in the measured dataset. (4) Remove directionality in ***C***. This step is required since metabolite fluxes can be affected either by upstream or downstream factors. (5) For the drug prioritization part, we remove STRING PPI edges from piTracer’s gene interaction network, to get a new network called ***G***. This step was introduced because STRING contains many hub genes, which could obfuscate relevant gene-to-metabolite connections by connecting virtually all genes to all metabolites in the piTracer network. (6) Combine ***G*** with ***C*** via metabolic genes (such as enzymes and metabolite transporters), which are present in both networks. (7) Calculate an inverse distance matrix ***Ꙇ*** between all genes in ***G*** and all metabolites in ***C*** in the ***G* + *C*** network. The inverse distance is calculated as *1/d*, where *d* is the number of steps between a gene-metabolite pair. ***Ꙇ*** has genes from ***G*** as rows and metabolites from ***C*** as columns. (8) Create a list ***E*** that contains genes directly interacting with metabolites, i.e., enzymes in metabolic reactions, in the ***G + C*** network. (9) To downweigh genes with many downstream metabolic target genes, assign each gene in ***G*** a weight defined as ***g*** = 1 - (# of ***E*** reachable)/|***E***|. The assumption here is that perturbing genes that regulate or interact with too many enzymes could have detrimental effects and focus on metabolic paths not relevant to the biological system. ***g*** downweighs these genes and will enable specific metabolic traces relevant to the system to be targeted. (10) Calculate gene scores ***S*** = diag(***g***)***Ꙇw***. For each gene, ***S*** contains the sum of the inverse distances *1/d* between the gene and all significant metabolites it reaches in the ***G* + *C*** network, weighted by the number of enzymes the gene reaches.

### 4.5 Cell culture and reagents

Cell line MDA-MB-231 (ATCC® HTB-26™) was obtained from the American Type Culture Collection (ATCC, Manassas, VA, USA) and propagated at 37 °C in a humidified 5% CO_2_ incubator. The cell line was cultured in Roswell Park Memorial Institute (RPMI)-1640 Medium, supplemented with D-(+)-Glucose, Sodium bicarbonate solution, 10% Fetal Bovine Serum Heat Inactivated (Sigma-Aldrich, MO, USA), and 1% Penicillin/Streptomycin (Thermo Fischer Scientific, MA, USA).

Glutaminase Inhibitor Compound 968 was purchased from Calbiochem (MA, USA). Imirestat and Ingenol Mebutate were purchased from MedChem Express (NJ, USA), Ranirestat from Sigma-Aldrich (MO, USA), and CAY10594 from Cayman Chemicals (MI, USA).

### 4.6 Cell viability assay

MDA-MB-231 cells were subjected to cell viability evaluation using 3-(4,5-dimethylthiazol-2-yl)- 2,5-diphenyltetrazolium bromide (MTT) colorimetric assay conducted in 3 separate experiments, each carried out in triplicates. NAD(P)H-dependent cellular oxidoreductase enzymes in living cells are capable of reducing the yellow tetrazolium dye MTT to a purple, water-insoluble MTT formazan, directly related to the presence of viable cells^58^. Cells were seeded at a density of 11,500 cells/well of a TC-treated 96-well microtiter plate. Cells were incubated in 37 °C in a humidified 5% CO_2_ incubator for 24h, prior to respective drug treatment.

For the single-agent study, MDA-MB-231 cells were treated with C968 (10 μM), Imirestat (C1 = 0.01, C2 = 0.1, C3 = 1, C4 = 10, and C5 = 100 μM), Ranirestat (C1 = 0.01, C2 = 0.1, C3 = 1, C4 = 10, C5 = 100 μM), CAY10594 (C1 = 0.001, C2 = 0.01, C3 = 0.1, C4 = 1, C5 = 10 μM), or Ingenol Mebutate (C1 = 0.01, C2 = 0.1, C3 = 1, C4 = 10, C5 = 100 μM).

For the combination experiments, MDA-MB-231 cells were treated with single agents: C968 (10 μM), Imirestat (10 μM and 100 μM), Ranirestat (10 μM and 100 μM), CAY10594 (1 μM and 10 μM), and Ingenol Mebutate (10 μM and 100 μM) and combination of agents: C968 (10 μM) with either Imirestat (10 μM and 100 μM), Ranirestat (10 μM and 100 μM), CAY10594 (1 μM and 10 μM), or Ingenol Mebutate (10 μM and 100 μM). All MTT experiments included control (untreated cells), and vehicle, cells treated with dimethylsulfoxide (DMSO). The cells were treated for 24h, 48h and 72h and the viability was monitored using MTT assay. Briefly, MTT (12 mM) solution in phosphate-buffered saline (PBS) was added into the cells following incubation at 37 °C. The supernatant was aspirated and the samples were frozen at −80 °C. The samples were thawed, DMSO (100 μL/well) was added following incubation at room temperature under constant shaking for 40 minutes. The absorbance was measured at 570 nm (reference wavelength 630) using CLARIOstar® microplate reader (BMG LABTECH). Absorbance values were used to calculate cell viability which was expressed relative to the value for DMSO-treated cells (controls, i.e., 100% viability). Significant differences in viability data were assessed via two-sample t-tests.

### 4.7 Cell lysis and Western blotting

For apoptosis monitoring using Western blot assay, cells were seeded in 6-well plates at a density of 0.48 × 10^6^ cells per well. The medium was changed 24h after seeding with a fresh 4 mL of conditioned media containing single agents: C968 (10 μM), Imirestat (10 μM and 100 μM), Ranirestat (10 μM and 100 μM), CAY10594 (1 μM and 10 μM), and Ingenol Mebutate (10 μM and 100 μM) and the combination of agents: C968 (μM) with either Imirestat (10 μM and 100 μM), Ranirestat (10 μM and 100 μM), CAY10594 (1 μM and 10 μM), or Ingenol Mebutate (10 μM and 100 μM). Cells treated for 24h with 1 μM staurosporine (Selleck Chemicals LLC) were used as a positive control for apoptosis detection. The cells were collected by trypsinization and the obtained cell pellets were further processed.

Whole cell lysates were prepared from the cell pellets in 2% SDS lysis buffer supplemented with protease–phosphatase cocktail inhibitors (Roche) and phenyl methane sulfonyl fluoride (Sigma-Aldrich). Samples were lysed by three freeze–thaw cycles in liquid nitrogen as described previously^59^. The supernatant was collected after centrifugation for 10 min at 12,000 rpm, and the total protein content was quantified using the DC protein assay (Bio-Rad, USA). The proteins were denatured by incubation with a 4× Laemmli buffer at 95 °C for 10 min.

The proteins prepared from whole cell lysates were used to conduct gel electrophoresis followed by transfer to a polyvinylidene fluoride membrane (PVD) (Bio-Rad, USA). The membrane was blocked in 5% milk solution in Tween-PBS (PBS with 0.1% Tween 20) for 1h and incubated overnight in a primary antibody at 4 °C. The overnight incubation was followed by three washing steps in Tween-PBS. The membrane was incubated in the corresponding secondary antibody followed by three washing steps in Tween-PBS. Both primary and secondary antibodies were prepared in the recommended dilution of either a 5% milk or 5% BSA solution in Tween-PBS. The signal was detected with a chemiluminescent Western blot detection kit (Thermofischer) and the blots were developed and visualized under a Bio-Rad ChemiDoc system (Amersham, Bio-Rad, USA).

The primary antibodies used were: Anti-AKR1A1 Antibody (RayBiotech, GA, USA), Aldose Reductase Antibody (H-6) (Santa Cruz, CA, USA), Anti-PLD2 antibody (Bio-Rad, CA, USA), Invitrogen SLC25A20 Polyclonal Antibody (Thermo Fischer, MA, USA), PARP (Cell Signaling, #9542), Caspase-3 (Cell Signaling, #9662), Caspase-7 (Cell Signaling, #9492), and B-tubulin (Cell Signaling, 2128S). The corresponding secondary antibodies included: horseradish peroxidase–conjugated anti-mouse (Cell Signaling, 7076S) and anti-rabbit (Cell Signaling, 7074S).

### 4.8 Cell death monitoring with high-throughput flow cytometry

MDA-MB-231 cells were treated for 48h with Vehicle (DMSO), C968 (10 μM), CAY10594 (10 μM) or Ingenol Mebutate (100 μM) alone or in combination of C968 (10 μM) withCAY10594 (10 μM) or with Ingenol Mebutate (100 μM). Cells treated with staurosporine (1 μM) for 24h were used as a positive apoptotic control. For apoptosis monitoring, cells were processed using FITC Annexin V Apoptosis Detection Kit according to manufacturer instructions (BD Biosciences, NJ, USA). Cells were harvested by trypsinization, washed twice with cold PBS and stained with fluorescein-conjugated Annexin V and PI according to the manual description (BD Biosciences, NJ, USA).

The percentage of cells undergoing apoptosis or necrosis was measured by flow cytometry using a LSR Fortessa from BD Bioscience equipped with a BD™ High Throughput Sampler (HTS). The high-throughput sampler was operated in Standard mode at the following settings: sample flow rate: 2.0 μl/s, sample volume: 100 μl, mixing volume: 100 μl, mixing speed: 200 μl, and four mixing cycles. Cell viability status was quantified based on the staining as follows: live (Annexin-FITC-negative, PI-negative), early apoptotic (Annexin-FITC+positive, PI−negative), late apoptotic (Annexin-FITC+ positive, PI+ positive) and necrotic (Annexin-FITC-negative, PI+positive) cells.

The measurements were performed in three independent experiments with three technical replicates each. Early and late apoptotic events were summed up to obtain an aggregate value, and technical replicates were averaged out. Statistical differences were assessed using two-sample t-tests between conditions across the three experiments.

## Acknowledgements

We thank the Flow Cytometry Facility within the Imaging Core at Weill Cornell Medicine - Qatar for contributing to these studies. The Core is supported by the “Biomedical Research Program at Weill Cornell Medicine - Qatar”, a program funded by Qatar Foundation.

