## Supplementary Materials for "piTracer - Automatic reconstruction of molecular cascades for the identification of synergistic drug targets"

*Supplementary Figure 1 – IPA and CPDB traces of central carbon metabolism and carnitine shuttle...2*
*Supplementary Figure 2 – TCA cycle ....... 3*
*Supplementary Figure 3 – ALDH18A1 to citrulline....... 4*
*Supplementary Figure 4 – MCCC1 and 3-hydroxyisovaleryl carnitine ....... 7*

#### Supplementary Figure 1 – IPA and CPDB traces of central carbon metabolism and carnitine shuttle

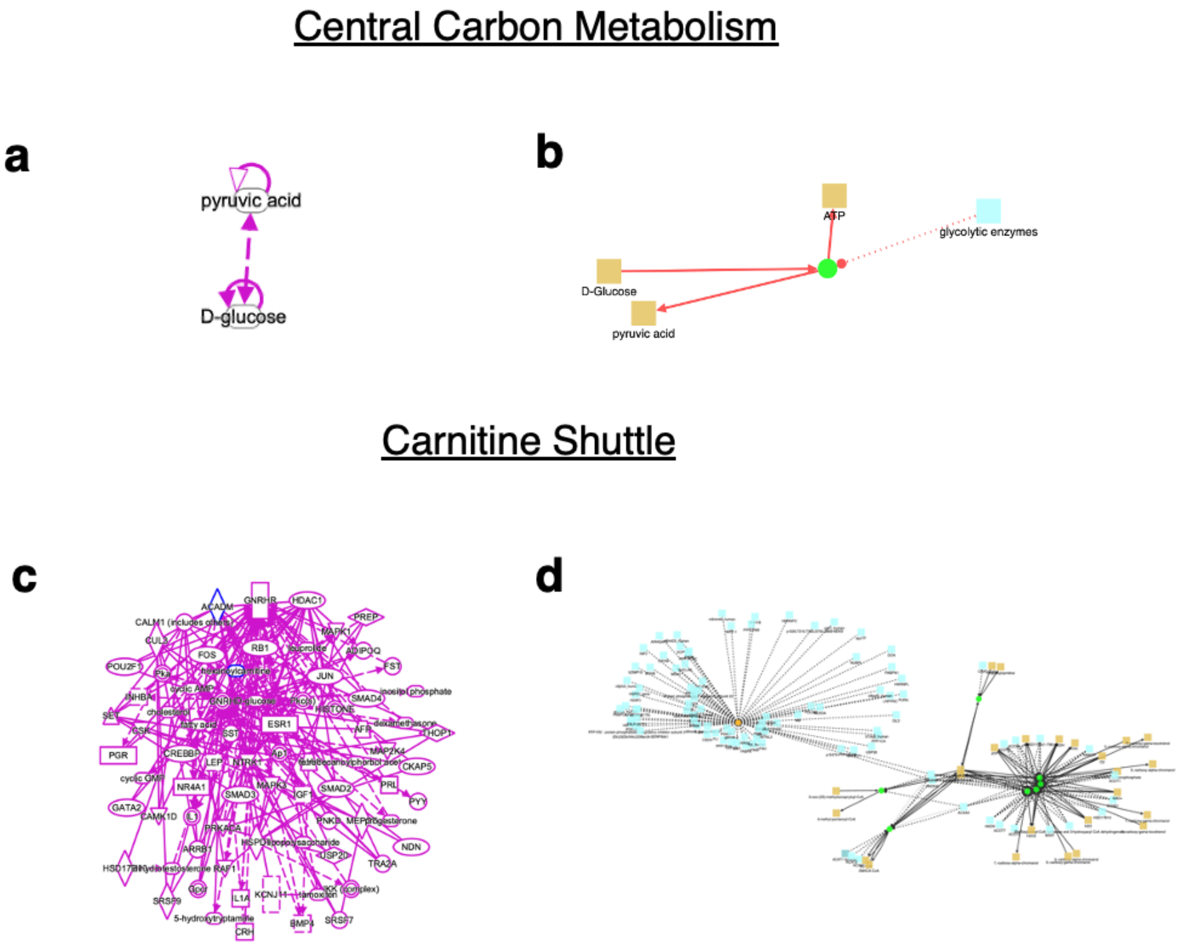

**Supplementary Figure 1.** (a) IPA and (b) CPDB reconstructions of central carbon metabolism between glucose and pyruvate. (c) IPA and (d) CPDB reconstructions of the carnitine shuttle between ACADM and hexanoyl carnitine. Note that ACADM and carnitine were used for the CPDB trace, since hexanoylcarnitine did not exist in its database.

To compare piTracer to existing pathway reconstruction tools, we calculated traces for central carbon metabolism as well as the carnitine shuttle mechanism in Ingenuity Pathway Analysis (IPA)[1] and ConsensusPathDB (CPDB)[2]. For central carbon metabolism between glucose and pyruvate, both IPA and CPDB simply connected the two molecules with no discernable path between them, which is probably caused by grouped pseudo-nodes in the underlying networks (**Supplementary Figure 1a&b**). For the carnitine shuttle between ACADM and hexanoylcarnitine, we had to use carnitine in CPDB, since hexanoylcarnitine did not exist in its database. Neither tool could recover the known mechanism connecting ACADM with hexanoylcarnitine. IPA recovered various paths that passed through gonadotropin-releasing hormone (GNRH), its receptor GNRHR, and a synthetic hormone leuprolide (**Supplementary Figure 1**c), while CPDB found paths between ACADM and carnitine through cofactors (**Supplementary Figure 1**d).

#### Supplementary Figure 2 – The citric acid cycle

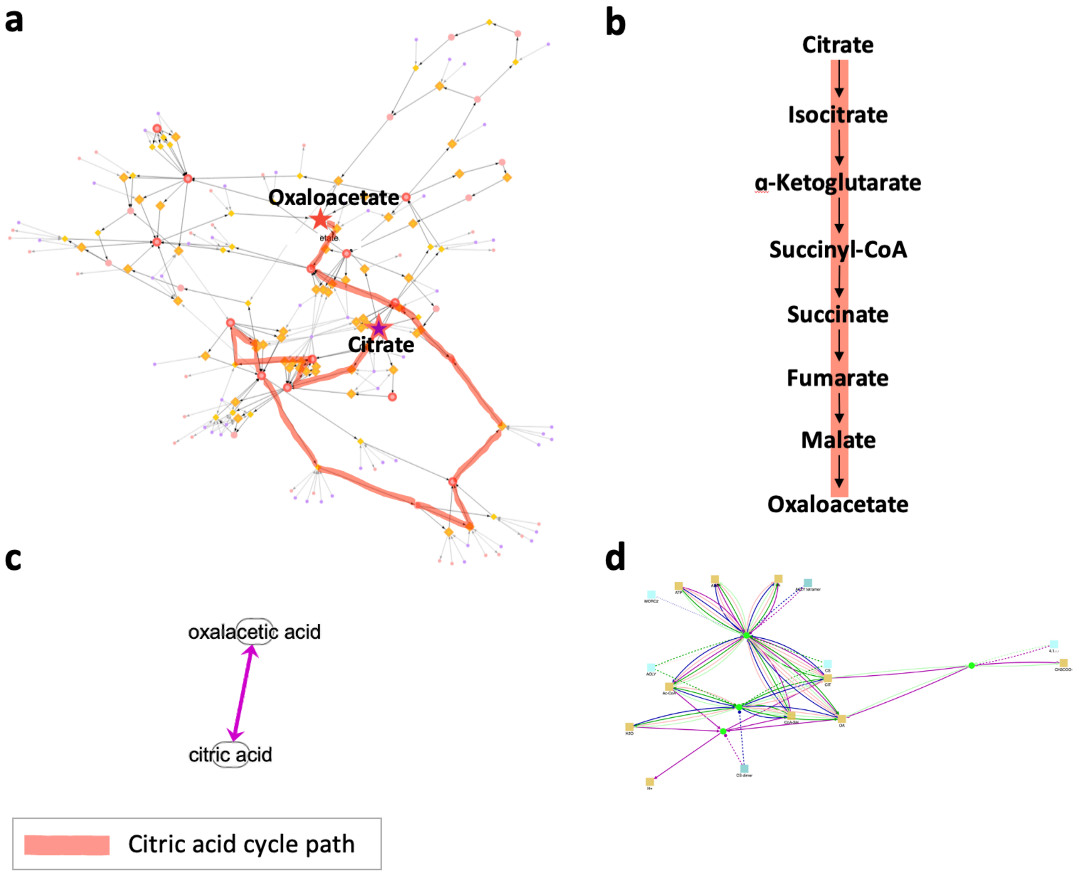

**Supplementary Figure 2.** Traces were produced by using citrate as the starting node and oxaloacetate as the end node. (a) Trace produced by piTracer. (b) Textbook version, (c) IPA trace, (d) CPDB trace of the pathway. Only piTracer was able to correctly reconstruct the citric acid cycle.

We further validated piTracer by tracing the top 10 shortest paths connecting citrate and oxaloacetate, the start and end point of the citric acid cycle. This pathway, also known as the tricarboxylic acid (TCA) cycle or the Krebs cycle, located in the matrix of the mitochondria in eukaryotes, is a core pathway for the metabolism of sugars, lipids, and amino acids[3]. By performing the piTracer query, we were able to find a path for the citric acid cycle (**Supplementary Figure 2a&b**) among other paths identified by piTracer. The TCA path also included a single reaction step between citrate and oxaloacetate. It must be noted that one biochemical step of the TCA, succinyl-CoA to succinate, was skipped in the trace, and α-KG was directly connected to the production of succinate instead. The reason for the absence of this biochemical step is that it did not exist in the multi-omics network used in the traces.

We compared our trace to the paths generated by IPA and CPDB. Both tools produced the same biochemically valid single reaction step between citrate and oxaloacetate (**Supplementary Figure 2c&d**). Neither CPDB nor IPA could reconstruct the citric acid cycle path. Notably, IPA contains a literature-curated version of the entire TCA pathway, but the automatic network reconstruction did not produce the correct results.

#### Supplementary Figure 3 – rs2403254 and 2-hydroxyisovalerate

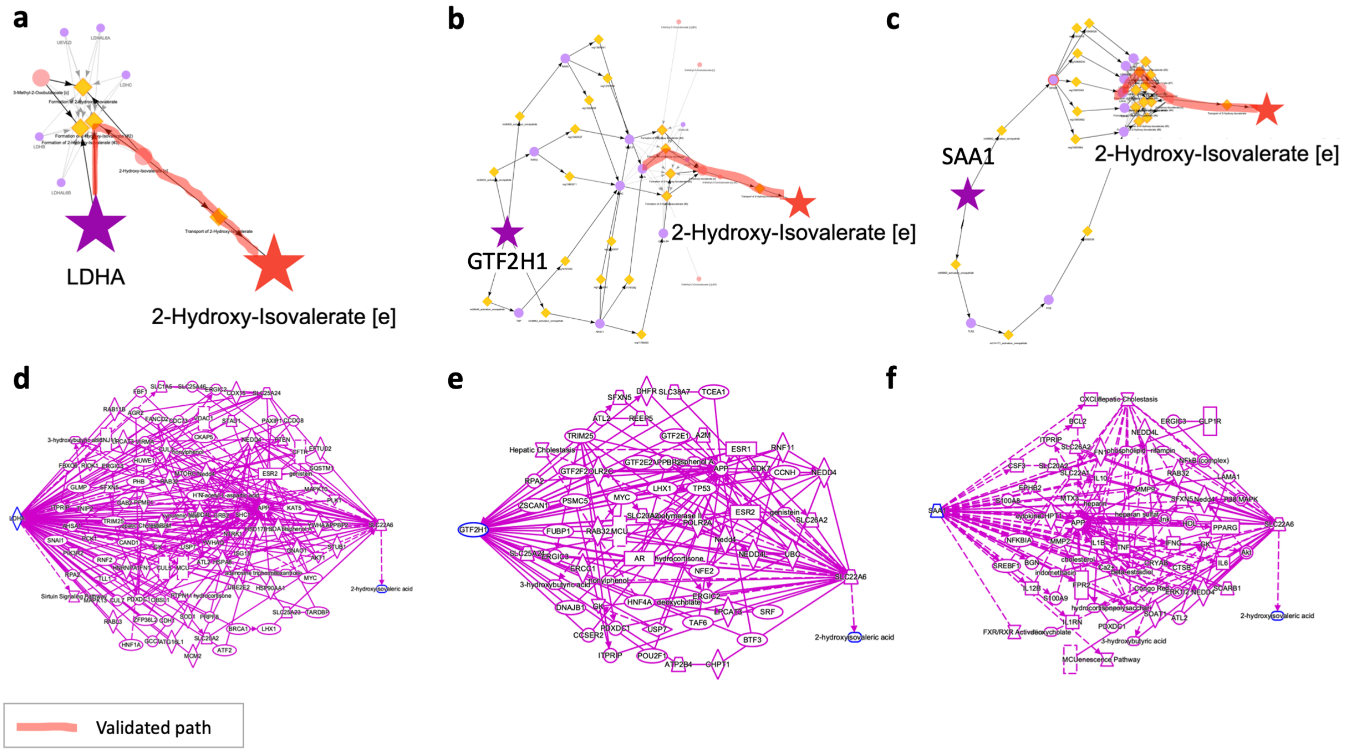

**Supplementary Figure 5.** Visualization of traces between (a) LDHA, (b) GTF2H1, and (c) SAA1 and 2-hydroxyisovalerate using piTracer. The highlighted paths in b and c are the same as the highlighted LDHA shortest path in a.

In their initial GWAS paper, for the association between SNP rs2403254 and 2-hydroxyisovalerate, Shin et al.[4] incorrectly annotated that locus with the HPS5 gene, since the SNP is located in in an intronic region of HPS5. Other candidate genes for the SNP found by a different study published by Heemskerk et al. were GTF2H1, SAA1, and LDHA[9]. The Heemskerk study demonstrated that LDHA was in fact the correct causal gene for the association, by showing that LDHA converts 3-methyl-2-oxobutanoate into 2-hydroxyisovalerate *in vitro*[9]. In contrast, the authors did not find any biochemical explanations for the HPS5-2-hydroxyisovalerate association. In the following, we assessed all four gene candidates using the piTracer framework.

First, there were no traces from HPS5 to 2-hydroxyisovalerate, since the gene does not have any functional annotations in our database, which is in line with the findings of Heemskerk et al. While we acknowledge that research bias might play a role here, the lack of functional information in extensive databases provides evidence against HPS5 being the correct gene. We then generated traces ($k$ = 10) from LDHA, GTF2H1, and SAA1 to 2-hydroxyisovalerate (**Supplementary Figure 5a-c**-) and subsequently ranked gene-to-metabolite associations by plausibility. We found that the LDHA trace contains the shortest path to 2hydroxyisovalerate (**Supplementary Figure 5-a**, highlighted path) and that LDHA directly catalyzes a reaction involving 2hydroxyisovalerate. Interestingly, the GTF2H1 and SAA1 traces, which consisted of longer paths, were extensions of the LDHA shortest path (**Supplementary Figure 5b&c**, highlighted paths). Using these traces, it would have been immediately clear that LDHA has the highest likelihood of being the correct gene associated with 2-hydroxyisovalerate.

We similarly traced between LDHA, GTF2H1, and SAA1 and 2-hydroxyisovalerate using IPA (**Supplementary Figure 5-d&f**). However, all traces had a constant path length, none contained the correct enzyme-metabolite relationship, and all were substantially longer and less specific than the shortest path found by piTracer. Additionally, we also found a trace between the wrong gene HPS5 and 2-hydroxyisovalerate with the same path lengths (not shown).

#### Supplementary Figure 4 – ALDH18A1 to citrulline

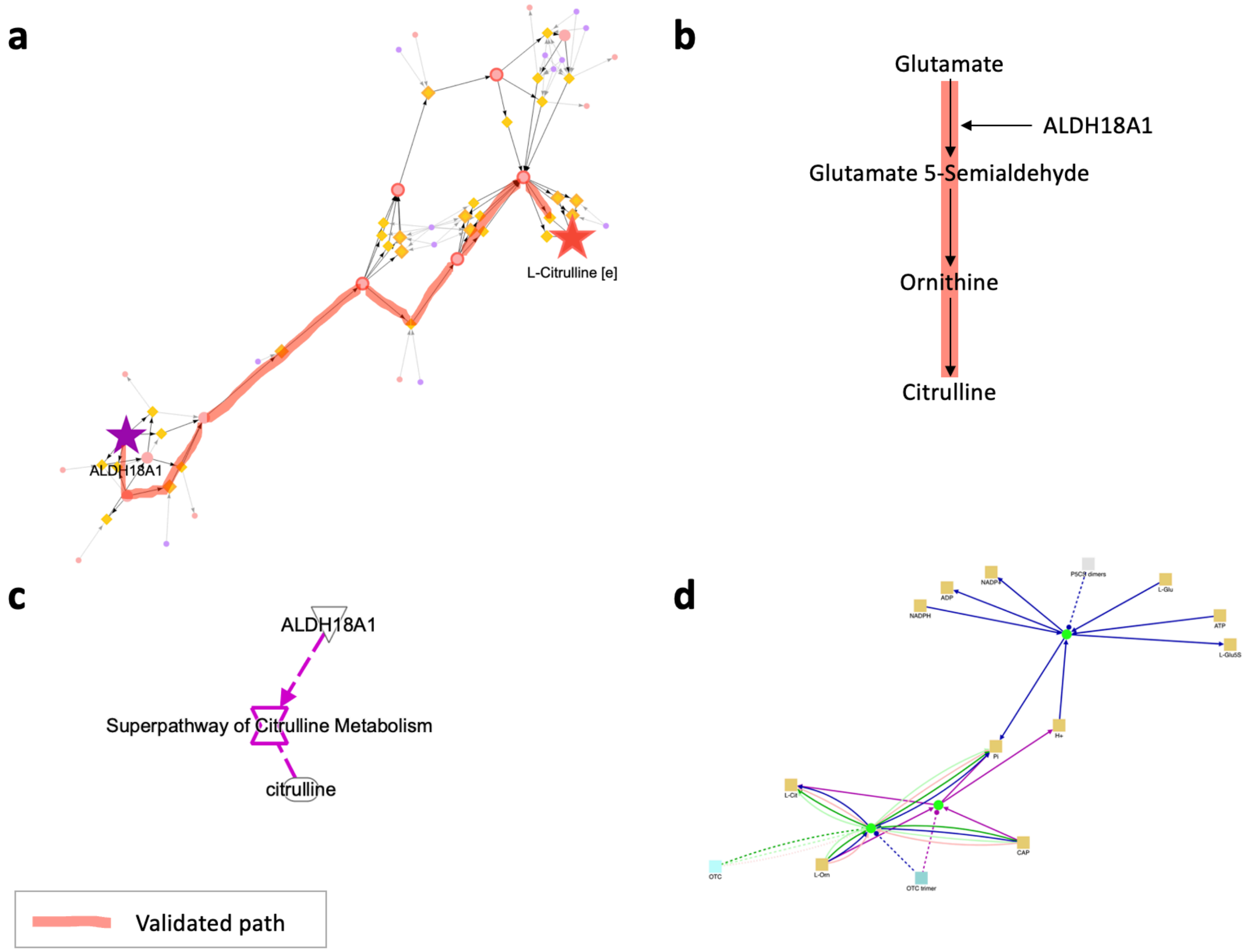

**Supplementary Figure 3.** (a) Trace produced by piTracer. (b) Textbook version, (c) IPA trace, and (d) CPDB trace of the pathway.

As another example, we traced between the ALDH18A1 gene and citrulline, an association described in Shin et al.[4]. By performing this query, we were able to reconstruct paths between ALDH18A1 and citrulline ( **Supplementary Figure 3a**). ALDH18A1 encodes delta-1-pyrroline-5-carboxylate synthetase, an enzyme that catalyzes a step in the *de novo* production of ornithine, which is subsequently converted to citrulline[5] (**Supplementary Figure 3b**). Mutations in ALDH18A1 can result in hypocitrullinemia[6] and other conditions. As illustrated (**Supplementary Figure 3a**, red highlighted path), piTracer successfully reconstructed a known biological cascade that connects ALDH18A1 and citrulline.

Furthermore, we traced between ALDH18A1 and citrulline using IPA and CPDB. However, IPA recovered a path that passed through a "super-pathway" (**Supplementary Figure 3c**). Although CPDB included ALDH18A1 and citrulline in the visualization, the gene and metabolite were connected via cofactors inorganic phosphate (Pi) and hydrogen ion (H+) (**Supplementary Figure 3d**). Similar to the TCA example above, IPA provides a manually generated version of this pathway, but the automatic and unbiased reconstruction does not work.

#### Supplementary Figure 5 – MCCC1 and 3-hydroxyisovaleryl carnitine

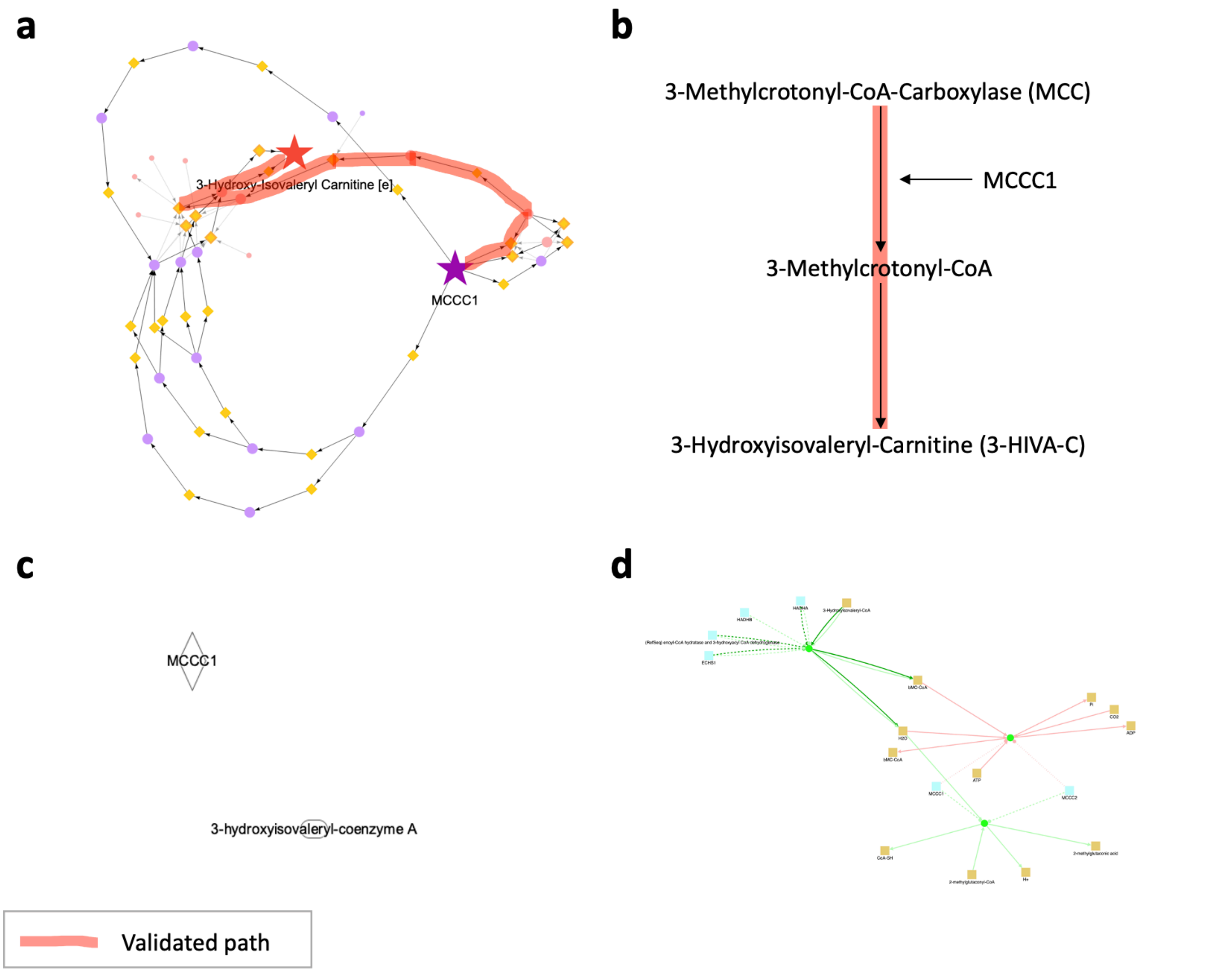

**Supplementary Figure 4.** (a) Trace produced by piTracer. (b) Textbook version, (c) IPA trace, (d) CPDB trace of the pathway. Note that for IPA and CPDB, MCCC1 was traced to 3-hydroxyisovaleryl-coenzyme A.

3-Methylcrotonoyl-CoA carboxylase 1 (MCCC1) catalyzes the conversion of 3-methylcrotonyl-CoA (3-MCA) to 3-methylglutaconyl-CoA in the breakdown of leucine. Alternatively, 3-MCA can be converted to 3-hydroxyisovaleryl-CoA (3-HICoA), a precursor of 3-hydroxyisovalerylcarnitine (3-HIC)[7]. MCCC deficiency has been associated with elevated plasma and urinary 3-HIC levels[8]. Moreover, Shin et al.[4] found that there was an association between MCCC1 and 3-HIC. We traced between the molecule pair (**Supplementary Figure 4a**) and found that the only metabolic path in the trace (red highlighted path) is in agreement.

Additionally, we attempted to construct traces between MCCC1 and 3-HIC using IPA and CPDB. Neither of the two tools yielded any paths. We further attempted to infer traces between MCCC1 and 3-HICoA instead (**Supplementary Figure 4c&d**), but only CPDB recovered a path from MCCC1 to 3-HICoA.

### Supplementary Figure 6 – Signaling pathways

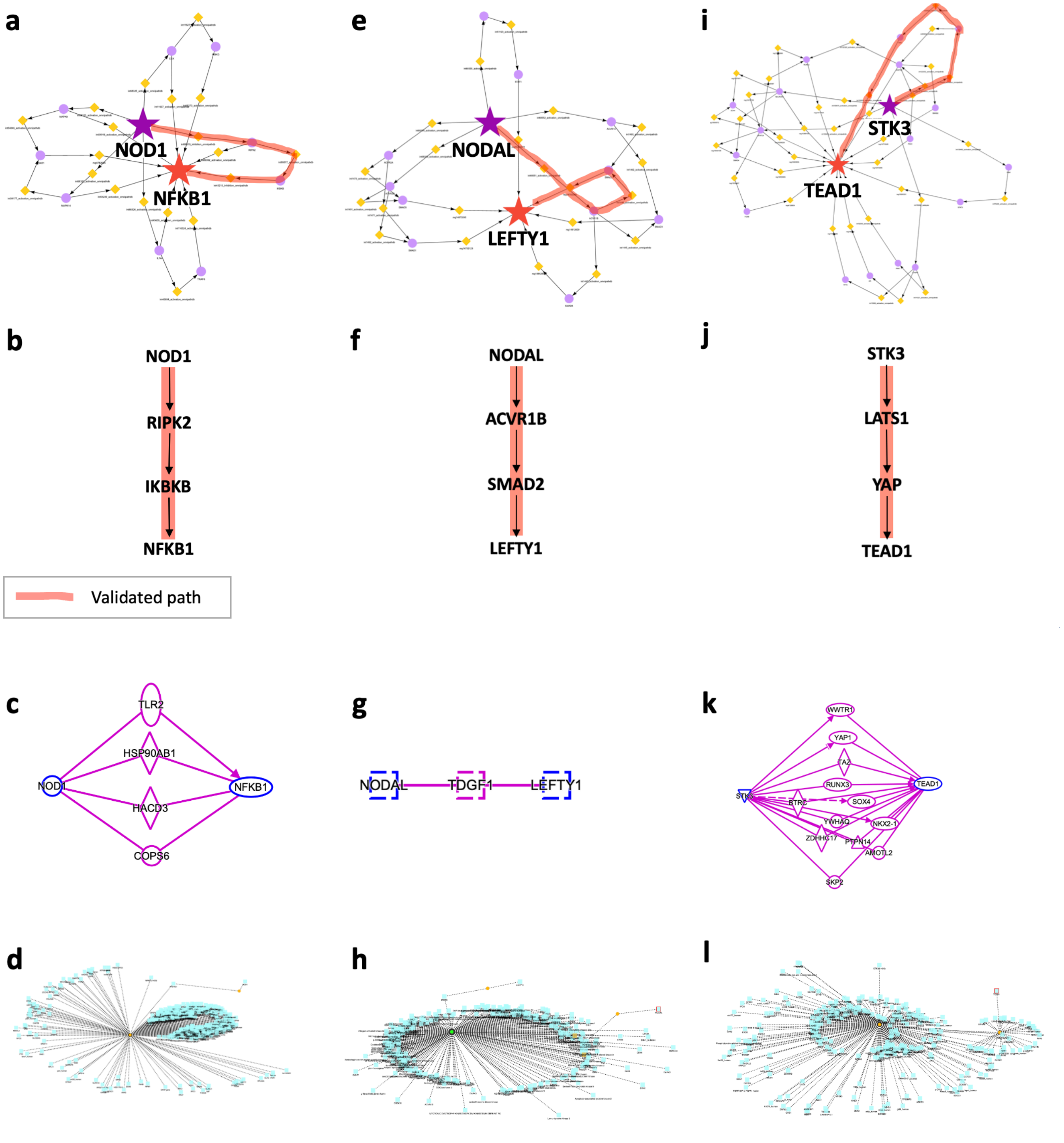

**Supplementary Figure 6.** piTracer Traces between (a) NOD1 and NFKB1, (b) known pathway, (c) IPA trace, and (d) CMPDB trace. (e) NODAL and LEFTY1, (f) known pathway, (g) IPA trace, and (h) CMPDB trace. (i) Hippo signaling pathway, (j) known pathway, (k) IPA trace, and (l) CMPDB trace.

##### *The NOD-like receptor signaling pathway*

We attempted to reconstruct the NOD-like receptor signaling pathway by tracing the top 10 shortest paths between NOD1 and NFKB1 (**Supplementary Figure 6a**). NOD1, together with NOD2, senses conserved motifs in bacterial peptidoglycan and induces anti-microbial responses and pro-inflammatory cytokines through NF-κB activation[10,11]. piTracer successfully found the major steps in the signaling pathway, namely the NOD1, RIPK2, IKBKB and NFKB1 path (**Supplementary Figure 6b**).

We also traced between NOD1 and NFKB1 using IPA and CPDB (**Supplementary Figure 6c&d**). Both IPA and CPDB indicated the existence of a connection between the two genes, however they did not capture any of the specific steps constituting the pathway shown in **Supplementary Figure 6b**.

##### *The NODAL signaling pathway*

As another example, we reconstructed the NODAL signaling pathway, by tracing between NODAL and its inhibitor LEFTY1 ($k$ = 10) (**Supplementary Figure 6e**). This pathway plays a central role in the maintenance of embryonic stem cell pluripotency, the patterning of the early embryo during mesoendoderm induction, and the dorsal-ventral axis specification in embryos[12]. Not only did our trace recover this signaling pathway ( **Supplementary Figure 6f**, highlighted path), but also included other equally valid paths that pass through different subcomponent combinations of Activin type 1 (ACVR1) and 2 receptors (ACVR2) and different SMADs involved in the pathway.

We also traced between NODAL and LEFTY1 using IPA and CPDB. While IPA connected the two genes in two steps using different genes (**Supplementary Figure 6g**), CPDB found a path between NODAL and LEFTY1 via MTOR, which is part of a different pathway, namely the mTOR signaling pathway (**Supplementary Figure 6h**).

##### *The Hippo signaling pathway*

As a third example, we reconstructed the Hippo signaling pathway using piTracer by tracing between STK3, a core kinase of that pathway, and TEAD1 (**Supplementary Figure 6i**). STK3 is a core kinase of the Hippo signaling pathway[13] and activates the transcription factor TEAD1 which leads to cell proliferation and survival[14], oncogenesis and chemotherapeutic resistance[15]. piTracer successfully reconstituted the pathway, cf. **Supplementary Figure 6j**. IPA included most major steps of the pathway, the LATS1/2 step (**Supplementary Figure 6k**). CPDB generated a trace that was not biologically meaningful (**Supplementary Figure 6l**).

#### Supplementary Figure 7 – Evaluation of abundance of selected protein targets under glutaminolysis inhibition.

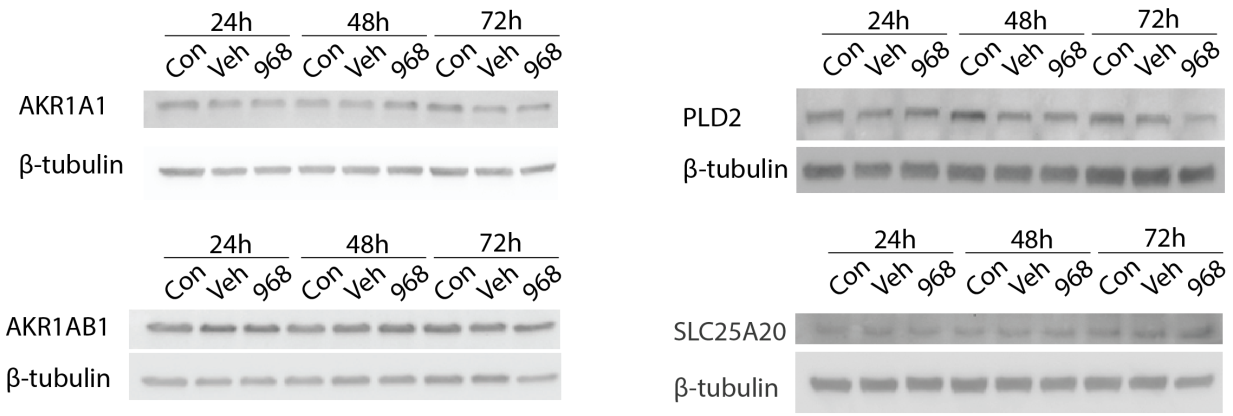

**Supplementary Figure 7.** Western blot analysis was conducted to evaluate whether C968 has an impact on the abundance of selected targets including: PLD2 (phospholipase D2); SLC25A20 (carnitine-acylcarnitine translocase, CACT); AKR1A1 (aldo-keto reductase family 1 member A1), and AKR1B1 (Aldo-keto reductase family 1, member B1). Lack of changes in the band intensities observed on Western blot suggests that treatment with C968 did not impact the expression of those protein. Cntrl - Control; Veh - vehicle; C968 - cells treated with 10 μM concentration of C968.

#### Supplementary Figure 8 – Viability of MDA-MB-231 cells after treatment with single drugs

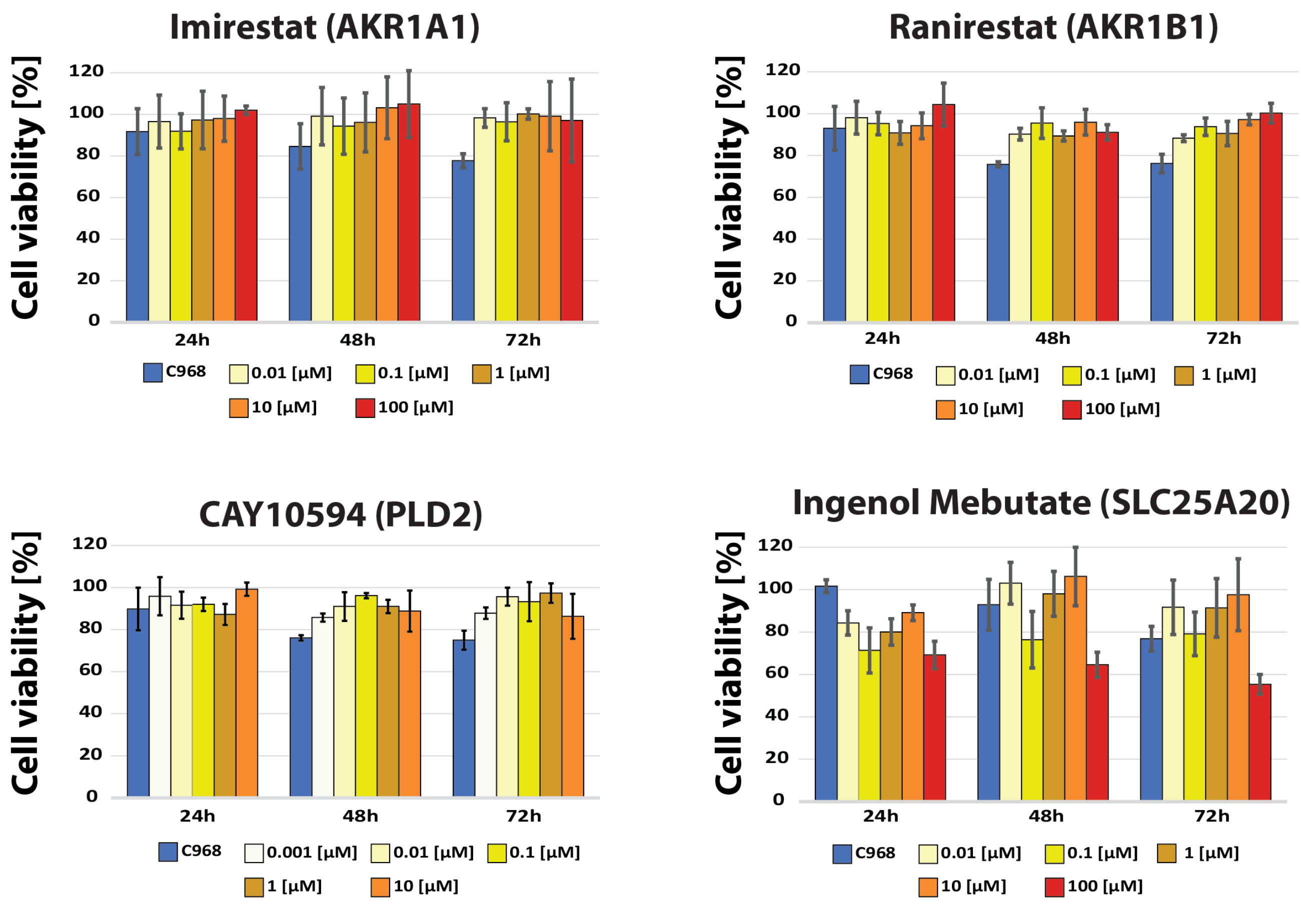

**Supplementary Figure 8.** The impact of selected inhibitors targeting AKR1A1/Imirestat, AKR1B1/Ranirestat, PLD2/CAY10594, and SLC25A20/Ingenol Mebutate, on the viability of MDA-MB-231 cells was tested using MTT assays. The components were tested at 5 different concentrations at three time points (24h, 48h and 72h). Overall, three out of four tested inhibitors had minimal impact on cell viability; this was observed for all examined concentration at all time points. A decrease in cell viability was observed when cells were treated with Ingenol Mebutate. The strongest ~40% decrease in cell viability was found at 72h after treatment with the highest concentration (100 μM) of Ingenol Mebutate. Cell viability was expressed relative to the value obtained for vehicle-treated cells (controls, i.e., 100% viability).

#### Supplementary Figure 9 – Impact of treatment strategies on the MDA-MB-231 cell line evaluated by microscopy

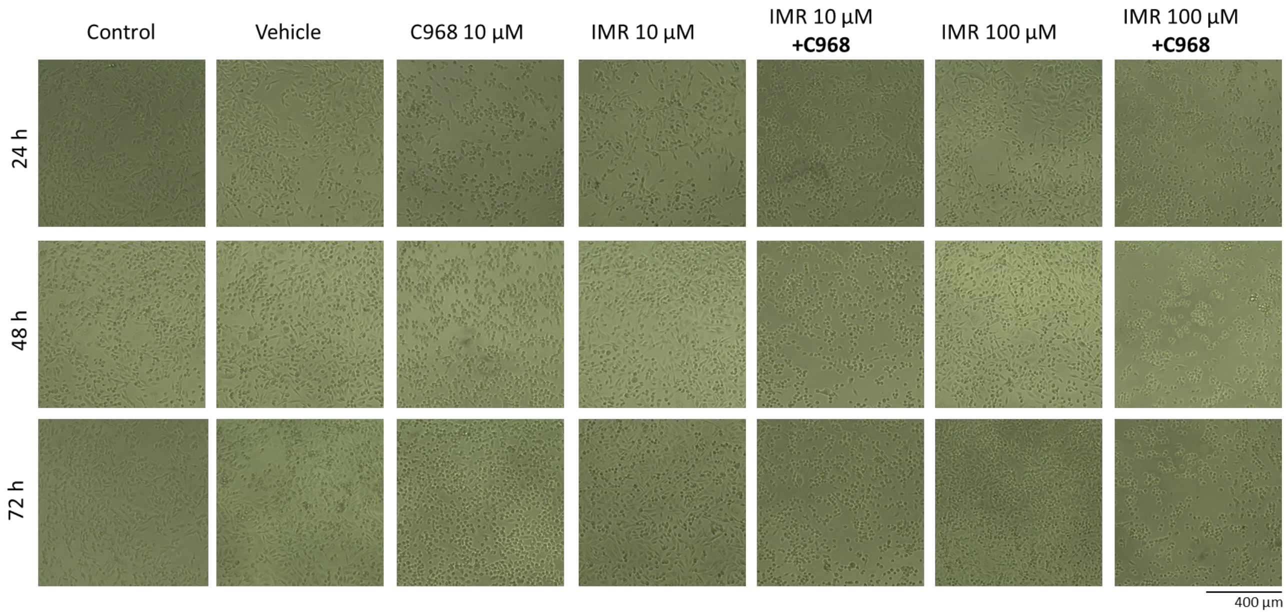

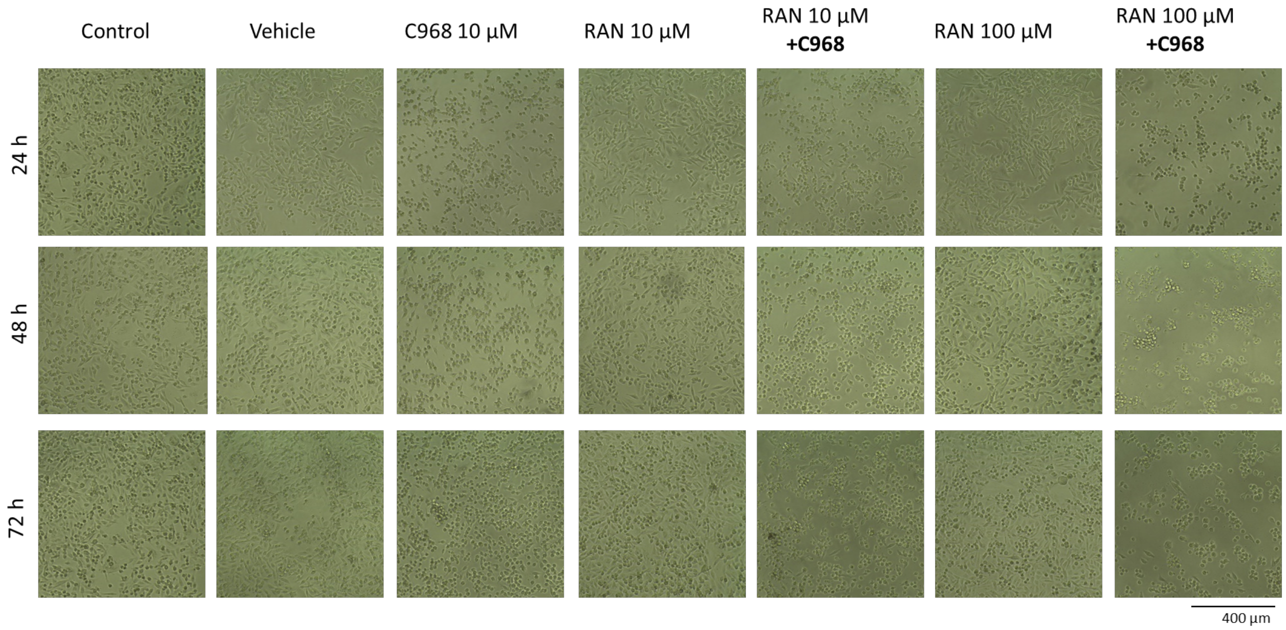

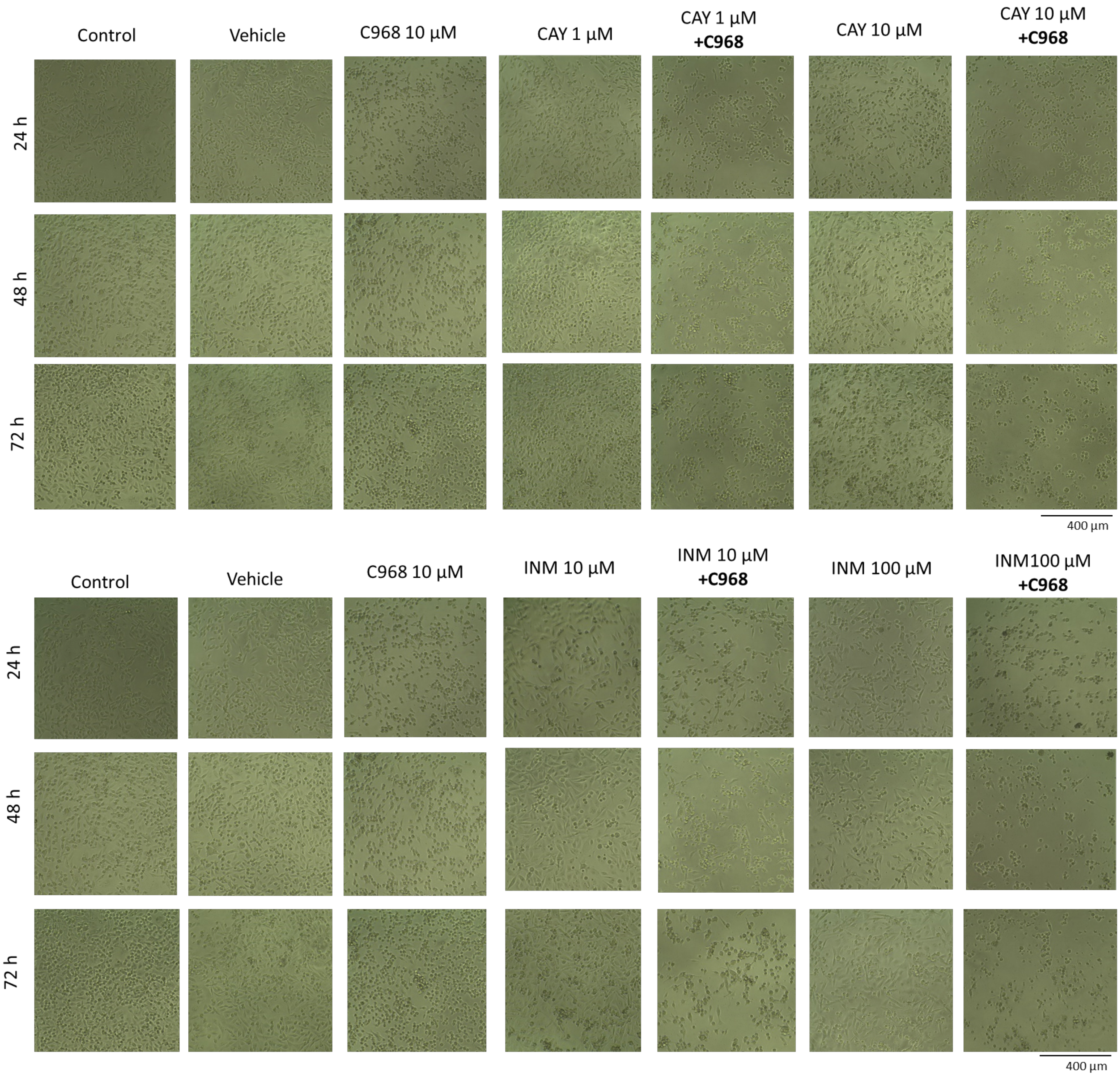

**Supplementary Figure 9**. The impact of single and combined treatment on MDA-MB-231 cells was evaluated at three time points (24h, 48h and 72h) using microscopy. Representative pictures are shown for each condition. As expected, there was an increase in cell density over time for both control and vehicle. Similarly, a time-dependent increase in density of cells treated with single drug agents was observed. A noticeable decrease in cell density was found for all combined treatment conditions. The most prominent effect was found at 72h after combined treatment with C968 and CAY (10 µM) as well as C968 and INM (100 µM). C968 – glutaminase inhibitor, IMR – Imirestat, RAN – Ranirestat, CAY – CAY10594, and INM – Ingenol Mebutate.

#### Supplementary Note 1 – Metabolic network scoring algorithm

To connect metabolites in our metabolic network in a pairwise fashion (metabolite-to-metabolite), we developed a specific scoring system that ensures meaningful links by tracing atoms from one molecule to the other. For a given molecule pair, we define two sets $M$ and $N$, which represent the respective sets of mapped atoms between them (intuitively, where to find the atoms of one molecule in the other molecule) according to the atom mapping algorithm in the Reaction Decoder Tool[16]. We then calculate the atomic overlap as $S_{MN} = |M \cup N|/|N|$. The overlap scores $S_{MN}$ and $S_{NM}$ are computed in a directional manner for all reactant/product and product/reactant pairs of each reaction.

However, simply applying an overlap threshold $T$ to $S_{MN}$ to determine which molecules should be connected in the metabolic network is not sufficient, because this would introduce shortcut paths or might miss important edges in the network. We illustrate this problem of simple thresholding in the following detailed example (see Figure below). Suppose that a given reaction 1 has reactant $A$ and products $B$, $C$, and $D$. Now assume that $S_{AB} = 0.02$, $S_{AC} = 0.49$, and $S_{AD} = 0.49$ in one direction, and $S_{BA} = 1$, $S_{CA} = 1$, and $S_{DA} = 1$ in the other direction. This means that 2% of atoms in $A$ are transferred to $B$, 49% of atoms in $A$ are transferred to $C$ and $D$, and reversely, all atoms of $B$, $C$, and $D$ are fully contained in $A$. Assume we set a base overlap score threshold $T$ of $0.75$. For this reaction, $S_{AB} < T$, but $S_{BA} > T$. A choice must be made on whether to create an edge between $A$ and $B$ in the network or not. If we require both the forward and backward overlap scores to be greater than $T$ ("AND" rule), then there will be no edges between $A$ and any of its products $B$, $C$, or $D$. Alternatively, we could introduce an "OR" rule that allows for the creation of an edge if either one of the overlap scores is $> T$. In that scenario, we would indeed have an edge $A$-$B$ ("Rule 1" in the Figure below).

Let us now inspect another reaction (Reaction 2 in the Figure below) with reactants $B$, $E$, and product $F$. Assume the overlap values are $S_{BF} = 1$, $S_{EF} = 1$, $S_{FB} = 0.01$, and $S_{FE} = 0.99$, where $B$ is the same molecule as in reaction 1. Note that $S_{FB} < T$ and $S_{BF} > T$. Given the "OR" rule, we create an edge $B$-$F$ ("Rule 1"). Inspecting both reactions 1 and 2 together, however, we now have a shortcut path $A$-$B$-$F$ in our network which is not biochemically reasonable, since $A$ and $F$ share no common atoms. In principle, $B$ acts as a cofactor in both reactions but does not contribute to the mass flow. If this procedure was applied to all reactions to construct a metabolic network, we would get a large number of shortcut paths passing through cofactors such as $B$. This would lead the metabolites from all reactions that use $B$ to be connected, creating various false positive edges and making the network unusable. In conclusion, neither the "AND" nor the "OR" rule generate satisfactory results.

To alleviate this problem, we instead calculate a reactant threshold value $T_m$ and a product threshold value $T_n$ defined as $T_m = T/N_m$ and $T_n = T/N_n$, where $N_m$ and $N_n$ represents the number of molecules on the other side of the reaction, i.e., the number of molecules the reactant is split into or synthesized from (see Figure below, overlap threshold table). We then check whether both $S_{MN} \geq T_M$ and $S_{NM} \geq T_N$. If both are greater than their corresponding thresholds, then an edge $M$-$N$ is created in the network ("Rule 2"). Intuitively, if a molecule splits into three products, then only a third of the atom overlap should be needed to define a reaction. Reversely, if a molecule is synthesized into a bigger molecule, it should be fully contained in the bigger molecule for this to be considered a metabolic edge. For our example, this means that we will exclude the shortcut edge $A$-$B$ as described in the following. Remember that $A$ splits into products $B$, $C$, and $D$, and thus $N_A = 3$. For reaction 1 and the pair $A$ and $B$, $T_A = \frac{T}{N_A} = \frac{0.75}{3} = 0.25$

and $T_B = \frac{T}{N_B} = \frac{0.75}{1} - 0.75$. Since $S_{BA} < T_B$ but $S_{AB} < T_A$, we will not create an edge $A$-$B$. If we calculate this rule for the other pairs in reaction 1, we find that we will create edges $A$-$C$ and $A$-$D$. For reaction 2 and the pair $B$ and $F$, we have $T_B = \frac{T}{T_B} = \frac{0.75}{1} = 0.75$ and $T_F = \frac{T}{N_F} = \frac{0.75}{2} = 0.375$. Since $S_{BF} \geq T_B$ but $S_{FB} \leq T_F$, we do not create an edge $B$-$F$. In contrast, we do create an edge $E$-$F$ following the same calculation. Now the shortcut path $A$-$B$-$F$ is effectively pruned from the network, leaving reaction 1 and reaction 2 correctly disconnected.

We used T = 0.75 for the processing of our metabolic network.

**Figure**

#### Reaction 1

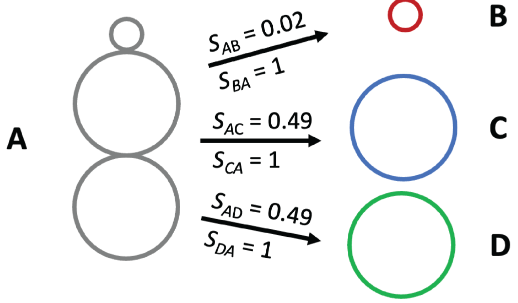

| Potential Edge | Edge? (Rule 1, "OR") | Edge? (Rule 2) |
|---|---|---|
| A –> B | Yes | No |
| A –> C | Yes | Yes |
| A –> D | Yes | Yes |

#### Reaction 2

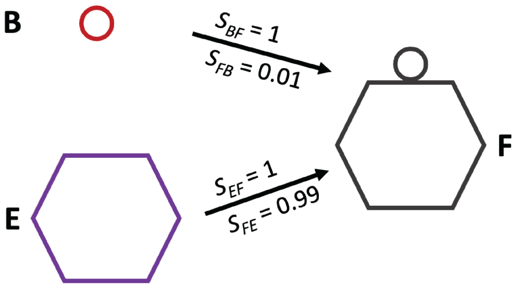

| Potential Edge | Edge? (Rule 1, "OR") | Edge? (Rule 2) |
|---|---|---|
| B –> F | Yes | No |
| E –> F | Yes | Yes |

#### Shortcut path

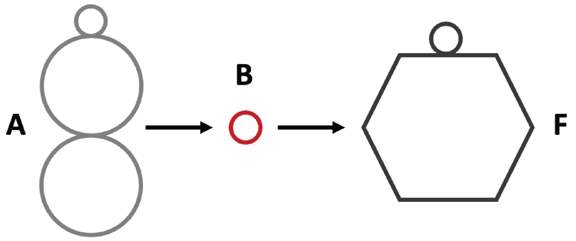

Given Rule 1, A –> B –> F shortcut exists
Given Rule 2, A –> B –> F shortcut is excluded

cofactor

**Overlap Threshold Values**

| Overlap score threshold | Split size | Why? |
|---|---|---|
| $T_A = T/N_A = T/3$ | $N_A = 3$ | A splits to B, C, D |
| $T_B = T/N_B = T/1$ | $N_B = 1$ | B is a product of A |
| $T_C = T/N_C = T/1$ | $N_C = 1$ | C is a product of A |
| $T_D = T/N_D = T/1$ | $N_D = 1$ | D is a product of A |
| $T_B = T/N_B = T/1$ | $N_B = 1$ | B produces F |
| $T_E = T/N_E = T/1$ | $N_E = 1$ | E produces F |
| $T_F = T/N_F = T/2$ | $N_F = 2$ | F is a product of B and E |
